# Cellular heterogeneity and therapeutic response profiling of human IDH+ glioma stem cell cultures

**DOI:** 10.1101/2025.07.29.667532

**Authors:** Nyasha Chambwe, Scott R Kennedy, Brendan F. Kohrn, Pavlo Lazarchuk, Mario Leutert, Guangrong Qin, Bahar Tercan, Monica Sanchez-Contreras, Weiliang Tang, Jerome J. Graber, Patrick J. Paddison, Judit Villén, Ilya Shmulevich, Raymond J. Monnat

## Abstract

Glioblastoma stem cell (GSC) cultures are initiated from glioblastoma (GBM) surgical resection tissue. They can capture and propagate key GBM primary tumor molecular and cellular features. We have deeply characterized four isocitrate dehydrogenase (IDH)-expressing (or IDH+) GSC cultures from unrelated adults to serve as cellular models for the majority of adult primary GBM. We demonstrate that GSC cultures can be continuously propagated in defined, serum-free media and 5% oxygen without requiring specialized growth substrates; have well-defined genomic and mtDNA variants and gene/protein expression profiles; and highly reproducible dose-survival curves when treated with the GBM standard-of-care therapies of ionizing radiation (IR) and temozolomide (TMZ). We also illustrate how expressed lentiviral barcodes, mtDNA variants and single cell gene expression profiling can be used to define and track cellular heterogeneity over 40 days after IR treatment. These well-characterized IDH+ GSC cultures can support many high throughput *in vitro* assay formats, including xenograft, organoid and other GBM disease modeling protocols. They should prove a useful resource to better understand GBM biology, and to identify new and more effective GBM therapies and treatment regimens.

**Highlights:**

- Glioblastoma (GBM)-derived IDH-expressing Glioma Stem Cell (GSC) cultures can capture and propagate GBM genomic variants, gene and protein expression programs.
- Mitochondrial DNA (mtDNA) variants identified by high accuracy Duplex DNA sequencing are abundant and sub-clonally organized in GSC cultures.
- GSC cultures have highly reproducible dose-survival curves for ionizing radiation and temozolomide, the GBM standard-of-care therapies.
- GSC culture cellular heterogeneity can be captured, characterized and tracked by using expressed lentiviral barcodes, mtDNA variants and scRNA sequencing.

**eTOC blurb:** IDH-expressing glioma stem cell cultures (GSCs) are experimentally tractable and versatile glioblastoma (GBM) cellular disease models. Deeply characterized GSC cultures can enable new work to understand GBM biology, and help identify new and more effective GBM therapies and treatment regimens.

**Graphical Abstract:** 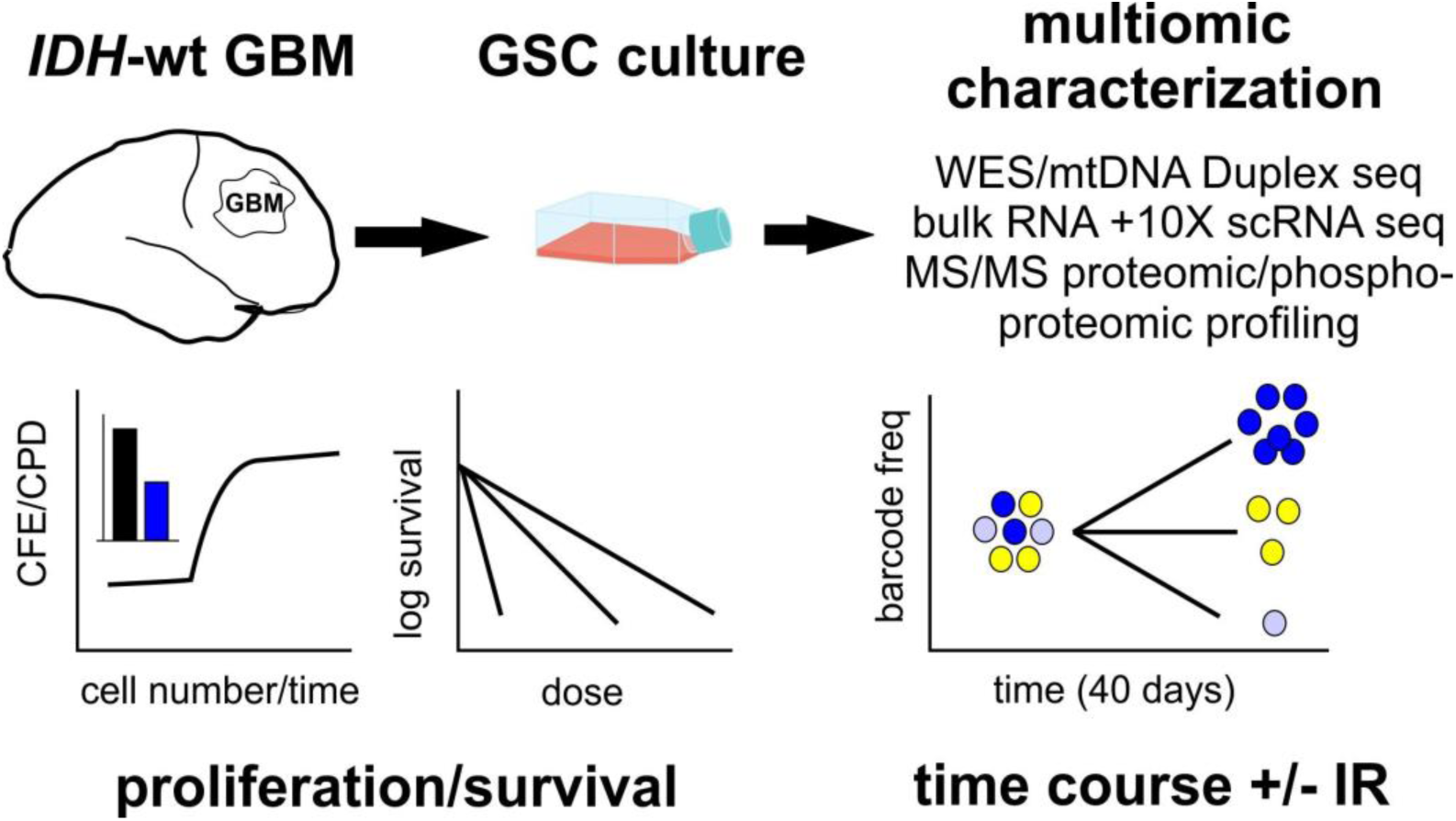

## Introduction

Gliomas are central nervous system tumors that originate from neuroglial stem or progenitor cells. The high histologic grade gliomas (Grade III anaplastic astrocytomas and Grade IV glioblastomas (GBM)) represent half of all primary adult brain malignancies, and are often fatal due to diffusely invasive growth and a lack of curative therapies (Louis et al., 2021; Schaff and Mellinghoff, 2023; Weller et al., 2024). Glioma genomic characterizations have identified key disease gene drivers and targets, including *IDH1* and *IDH2* neomorphic mutations and disrupted kinase signaling networks (Brennan et al., 2013; Cancer Genome Atlas Research Network, 2008; Parsons et al., 2008) (Liu et al., 2020; Yan et al., 2009). These genomic data have substantially improved glioma classification and prognosis, but have not as yet led to more effective therapies or improved patient outcomes (Louis et al., 2021; Schaff and Mellinghoff, 2023; Weller et al., 2024)

Work to better understand GBM biology and therapeutic vulnerabilities continue to depend on tractable and revealing preclinical disease models (Cirigliano and Fine, 2025). Well-characterized glioma stem cell (GSC) cultures address this need: they are initiated directly from GBM surgical resection tissue; are enriched in tumor stem/repopulating cells, a key therapeutic target; capture and can propagate many GBM molecular and phenotypic features; and are experimentally more versatile, cheaper and easier to use than many other GBM disease models. GSC cultures can also support a growing array of high-throughput *in vitro* protocols as well as increasingly useful organoid, xenograft and on-chip disease modeling protocols (Cirigliano and Fine, 2025; Furnari et al., 2024; Gimple et al., 2019; Lathia et al., 2015; Suvà and Tirosh, 2020).

We have systematically characterized four isocitrate dehydrogenase 1/2-expressing (IDH+) GSC cultures that are representative of ∼90% of newly diagnosed IDH+ adult primary GBM (Weller et al., 2024). All four GSC cultures were initiated from surgical resection tissue from unrelated adults (Son et al., 2009; Wakimoto et al., 2009); represent previously defined GBM genomic, gene expression and proteomic subtypes; and have reproducible dose-survival curves for GBM standard-of-care therapies ionizing radiation (IR) and temozolomide (TMZ). Two types of DNA-based molecular barcodes and single cell gene expression profiling were used to define GSC cell/state heterogeneity, and to track single cell trajectories over 40 days after IR treatment. These well-characterized, experimentally tractable GSC cultures should help improve our understanding of GBM biology, and better enable the search for new and more effective GBM therapies and treatment regimens.

## Results

### Propagation, growth rate and colony-forming efficiency

GSC cultures GBM4, GBM8, 0131 and 0827 were initiated from surgical resection tissue from unrelated, anonymized adult GBM patients in defined, serum-free media. Three cultures were from primary GBM resection tissue; GSC culture 0827, as we document below, was found to have originated from the recurrence of a previously IR+TMZ-treated primary GBM (Son et al., 2009; Wakimoto et al., 2009). All four cultures were propagated in defined, serum-free media at 37°C in a 5% CO_2_/5% O_2_ atmosphere: specialized growth substrates were not required, though all four cultures retained the ability to grow as monolayers when plated on laminin-coated plastic (Toledo et al., 2015). Prior to use, all cultures were authenticated by short tandem repeat (STR) DNA fingerprinting, and verified to be *Mycoplasma*-free using a common workflow (**Figure 1A, Table S1**).

**Figure 1:**
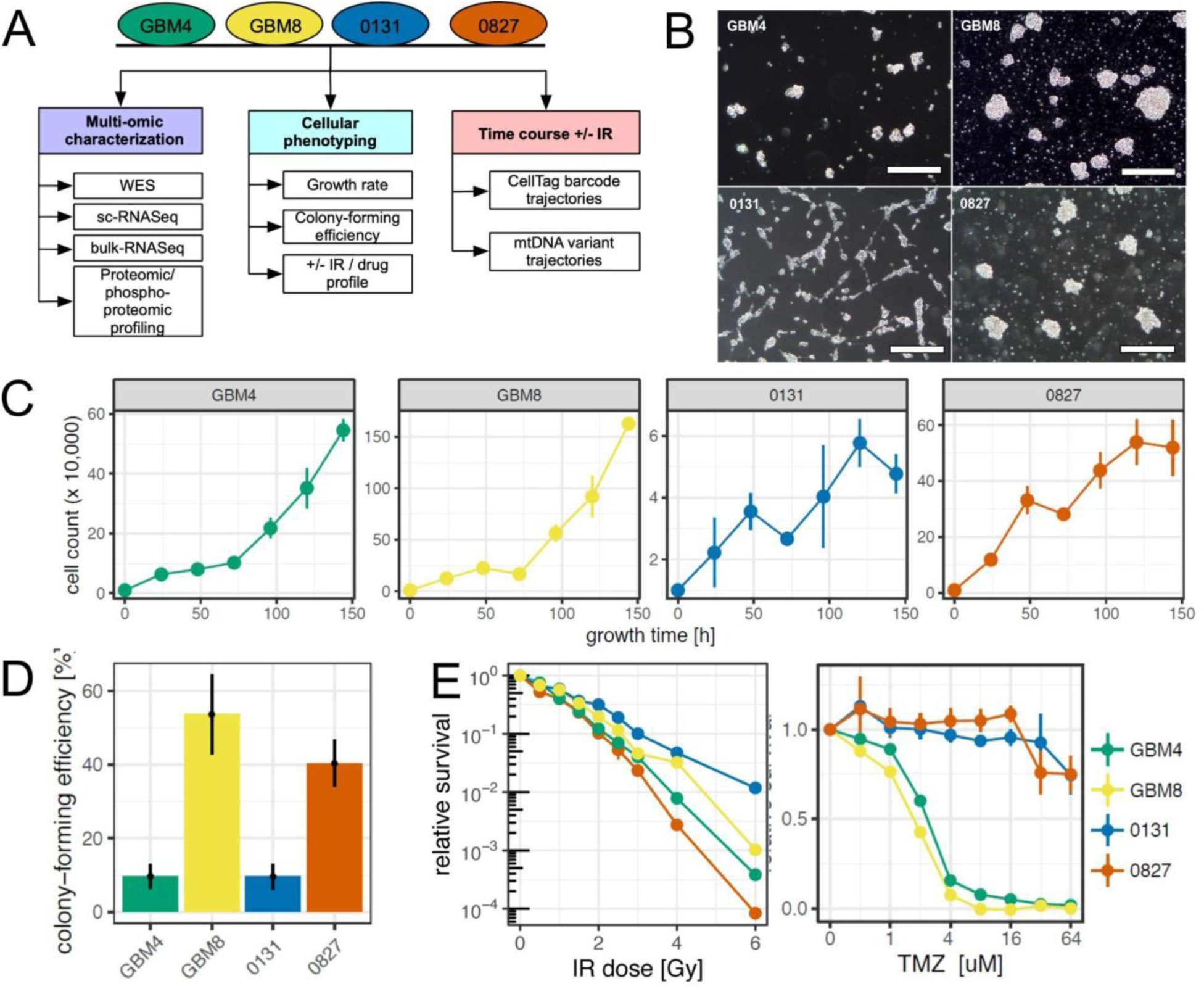
G**l**ioma **stem cell (GSC) proliferation and therapeutic response profiles.** (**A**) GSC culture experimental workflow. (**B**) Dark field microscopy of GSC cultures in mid-exponential growth in serum-free medium, 5% oxygen on untreated plastic. Scale bars are 200 μm long. (**C**) GSC proliferation kinetics reveals similar trajectories but substantially different population doubling times (note Y-axis units for cell number). (**D**) Colony-forming efficiencies (CFE) of untreated GSCs determined by dilution cloning. (**E**) Heterogeneity of dose-response as a function of IR or TMZ dose versus untreated controls quantified by CFE. ***Abbreviations:*** CellTag, lentiviral molecular barcoding system (Biddy et al., 2018); IR, ionizing radiation; mtDNA, mitochondrial DNA; TMZ, temozolomide; scRNAseq, single cell RNA sequencing; WES, whole exome sequencing. Error bars represent standard error of the mean in replicate experiments.

GBM4, GBM8 and 0827 cultures grew in suspension from dispersed single cells to form multicellular neurosphere-or tumoroid-like aggregates. Culture 0131, in contrast, grew from substrate-attached single cells to form eventual attached and suspension aggregates (**Figure 1B**). Bulk population doubling times of GBM4, GBM8 and 0827 were ∼ 24 hrs, whereas 0131 had a substantially slower doubling time (**Figure 1C**). All cultures readily generated single cell-derived colonies by dilution cloning, with colony-forming efficiencies of from 10% to 53% (**Figure 1D**).

### Radiation and drug dose-survival analyses

Cell survival as a function of dose was determined using both bulk population growth suppression and colony-forming efficiency (CFE) assays. IR doses of 1.0 - 1.5 Gy reduced survival by ∼67%, with culture-specific CFE differences ranging up to ∼100-fold at 6 Gy (**Figure 1E, left panel**), where IR-treated survival was not well-correlated with either growth rate or growth fraction (**Figures 1C, E and S1**). All four cultures could however be IR-sensitized by pre-treatment with the ATM kinase inhibitors KU-60019 or AZD-1390, leading to 3-to ∼40-fold reductions in CFE as a function of IR dose. Only GBM4 displayed ATM inhibitor-dependent CFE suppression in the absence of IR (**Figure S1A**).

Cultures GBM4 and GBM8 displayed steep dose-dependent TMZ survival curves with LD_50_ values of 2 - 3 µM, whereas 0131 and 0827 were effectively resistant at all TMZ concentrations tested (to 64 µM in **Figure 1E**, and to 250 uM in additional data not shown). Only 0131 expressed methyl-guanine methyltransferase (MGMT) protein, a major determinant of TMZ resistance, as documented by RNAseq and Western blot analyses (Hegi et al., 2005; Stupp et al., 2005)(**Figure S1B**). In order to better assess the role of MGMT expression in TMZ resistance, we grew cultures in the demethylating agent decitabine to induce MGMT expression. However, all four cultures were killed at sub-micromolar decitabine concentrations, leaving too few surviving cells to permit MGMT expression analyses (**Figure S1C**).

All four GSC cultures were also killed by low nM concentrations of ST-401, a small microtubule-targeting molecule already known to be effective in targeting mouse GBM xenografts that kills cells by a mechanism distinct from taxol or vincristine (Horne et al., 2021; Vicente et al., 2024)(**Figure S1D**). Pilot experiments with GBM4 were performed to determine whether exogenous glutamate altered GSC viability or proliferation (Iacovelli et al., 2018; Venkataramani et al., 2019). However, there was no effect on proliferation or survival up to concentrations of 2 mM after controlling for glutamate addition-associated pH effects (data not shown).

### Cytogenetic characterization

All four GSC cultures were aneuploid in metaphase cytogenetic analyses with numerous numerical and structural abnormalities and mean chromosome numbers ranging from 51 - 86. GBM4 and GBM8 were near-tetraploid, whereas 0827 was near-triploid (**Figure S2**). X chromosome gains or losses were present in 3 cultures, and all four displayed previously reported, recurrent GBM chromosomal abnormalities: partial or complete chromosome 7 gain (GBM8 and 0131); chromosome 20 gain; and chromosome 4 and 10 loss (GBM4 and GBM8). Additional abnormalities included marker chromosomes, and ‘double minute’ chromosomes in GBM4 and GBM8 (**Figure S2)**(Bigner et al., 1988; Dahlback et al., 2009).

### Genomic variant identification

We used whole exome sequencing (WES) to identify GSC single nucleotide variants (SNVs) and small insertion/deletion (indel) variants of ≤10 bp. GSC-specific variants were then compared with comparable data on 2,539 primary GBM tumor samples identified in cBioportal (https://www.cbioportal.org/) as ‘CNS/Brain - Cancer Subtype Designation’‘Glioblastoma multiforme’ samples (accessed 20250407). These comparison samples were almost exclusively (97%) primary GBM tumors, among which we identified 212 genes by OncoKB criteria that were adequately sampled (≥ 10 samples/2,539 tumors) and with altered frequencies of >1%. Twenty-two of these genes harbored 130 variants with from 22 to 54 variants/culture: there were from 3 (GBM8) to 36 (0827) variants/GSC culture among the top 25 genes altered by frequency in our GBM reference set (**Figure 2A, Table S2**).

**Figure 2:**
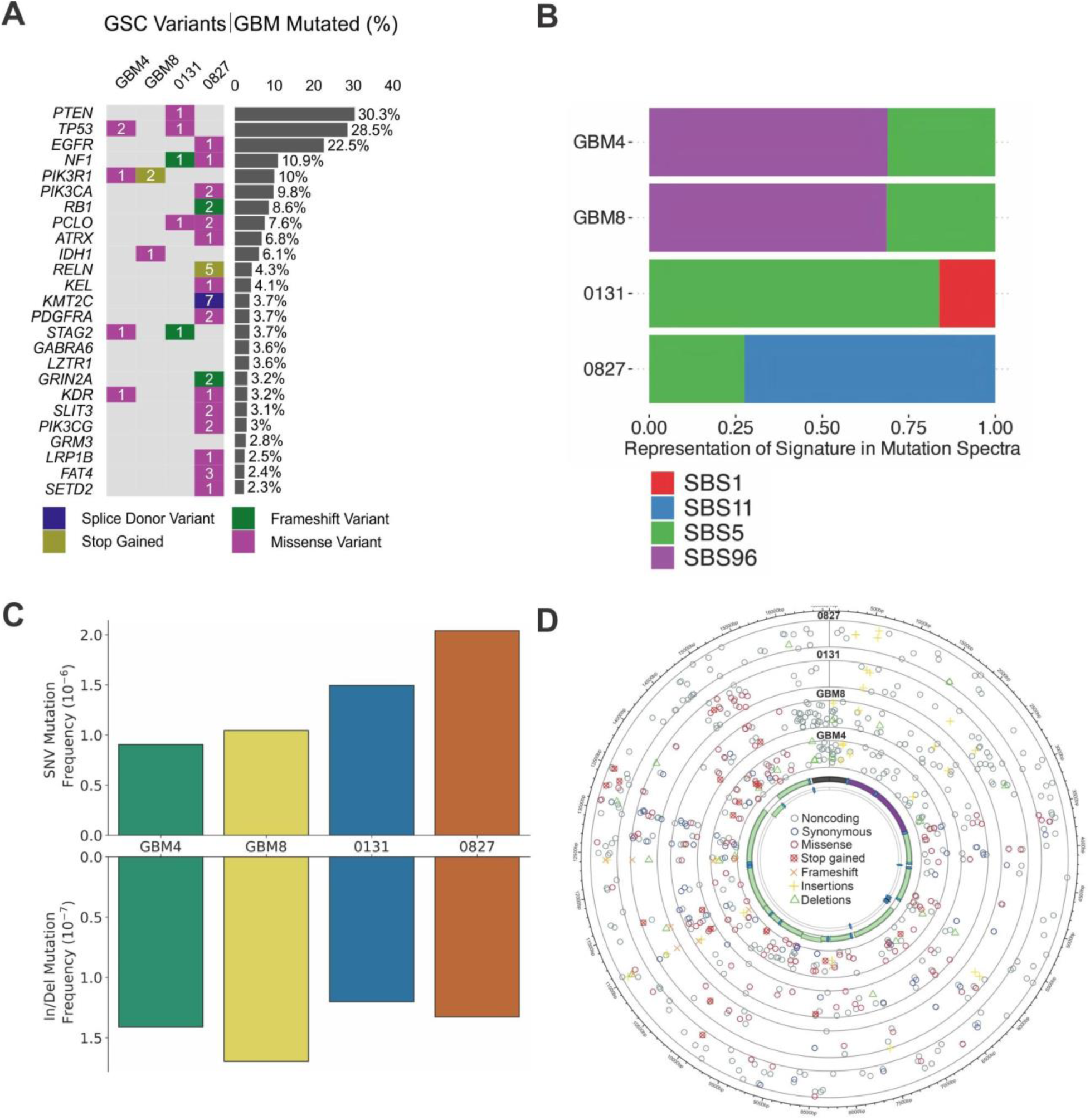
G**S**C **culture genomic and mtDNA variants identified by whole exome and Duplex mtDNA sequencing. (A)** GSC variants identified by whole exome sequencing in the top 25 genes altered in a CBioportal GBM reference set of 2,593 samples. Numbers in cells indicate GSC variants/gene, with the cell color code indicating the worst predicted consequence of GSC/gene-specific variants. **(B)** Barplot of fractional representation of GSC culture single base substitution (SBS) mutational signatures among top-ranked COSMIC SBS signatures. GBM4 and GBM8 had virtually identical mutation distributions attributable to SBS5 (446 and 445 mutations, respectively) and to SBS96 (987 and 972 mutations, respectively)(see Methods for detail). **(C)** mtDNA single nucleotide (SNV) and insertion/deletion (indel) variant frequencies in GSC cultures. Note Y-axis difference in SNV versus indel units. **(D)** Circos plot of mtDNA variant locations, molecular types and predicted functional consequences in GSC cultures displayed on human mtDNA complementary strands (center) with coding regions shown for proteins (*light green*), rRNAs (*purple*) and tRNAs (*blue*).

Most of the 130 genomic variants identified by WES, 88.5%, were missense variants with little or no predicted functional impact despite being located in genes previously identified as part of key GBM receptor tyrosine kinase (RTK), RAS and PI3K signaling pathways where deleterious somatic or germline variants are known to influence glioma risk or pathogenesis (Choi et al., 2023; Eckel-Passow et al., 2022; Graber, 2025; Savage et al., 2025). No GSC *IDH1* or *IDH2* neomorphic or *MGMT*-inactivating variants were identified, though we did identify several previously reported non-synonymous SNVs and indels in 0131, 0827 and GBM8 (Toledo et al., 2015)(Wang et al., 2018b). Fifteen predicted pathogenic variants were identified by a combination of variant molecular type, predicted biochemical consequence, and high variant effect prediction scores using REVEL (Ioannidis et al., 2016) or AlphaMissense (Cheng et al., 2023) pathogenicity scoring. These variants induced frameshifts and/or truncated open reading frames (n=11); induced mis-splicing (n=1); or led to internal duplication-deletions (n=3)(see Methods for scoring details)(**Table S2**). Potential clinical actionability of these predicted deleterious variants was assessed using the ‘OncoKB Actionable Genes - Glioma’ resource (www.oncokb.org/actionable-genes#cancerType=Glioma&sections=Tx; update API 27 February 2025) and the COSMIC ‘Mutation Actionability in Precision Oncology’ resource (version 16, 21 May 2025, see: cancer.sanger.ac.uk/cosmic/download/actionability): eleven altered ‘Top25’ genes in GSC cultures were potentially actionable in CNS gliomas and related neoplasms using existing approved drugs, with additional drugs in trials or with strong biological rationales for therapeutic development (**Table S2**).

Exome sequencing data also allowed us to estimate GSC-specific tumor mutational burden (TMB), an important immune oncologic (IO)-targeting metric. Aggregate median GSC culture TMB was 3.76 mutations/Mb (range 3.36 - 67.86 mutations/Mb), resembling the median TMB estimate of 2.6/Mb for 10,294 glioma samples (Touat et al., 2020), and 1.04/Mb for a TCGA cohort of 359 *IDH*wt samples. GSC culture 0827 was a clear TMB outlier with a median TMB of >50.8, or closely resembling reported hypermutated *IDH*-wildtype GBM samples. We did not identify an obvious genomic driver of this high 0827 TMB, e.g., mismatch repair loss and/or the presence of *POLD1/E1* mutator alleles (Touat et al., 2020) (Cheng et al., 2024; Haynes et al., 2024; Richardson et al., 2023). However a 0827 mutational signatures analysis revealed a strong SBS11 (Single Base Substitution) and SBS5 signatures representing, respectively, 72.5% and 27.5% of the 0827 signature. The dominant SBS11 signature has been attributed to TMZ DNA base damage, and the SBS5 signature to an as-yet unidentified clock-like source (per SigProfiler - Human Cancer version 3.4, accessioned 20250414: (Shen et al., 2019)(Brennan et al., 2013)(**Figure 2B**). Consistent with these results, we subsequently learned that 0827 originated from a recurrent or secondary GBM, where the primary had been treated 4 years earlier with IR and TMZ (personal communications, Patrick Paddison, Jeongwu Lee (Cleveland Clinic Center for Cancer Stem Cell Research) and P.J. Cimino (Neuropathology Head, NIH Surgical Neurology Branch)). Thus 0827 TMB, cellular and molecular features need to be interpreted in light of this history of prior treatment, with potentially strong selection during regrowth.

### Mitochondrial DNA variation in GSC cultures

High-accuracy Duplex DNA Sequencing identified 479 unique mtDNA variants, and allowed us to further define their subclonal architecture in GSC cultures. Most variants (∼90%) were single nucleotide substitutions (SNVs), with from 74 (0131) to 122 (GBM4) unique variants/GSC culture. Variant Allele Fractions (VAFs) ranged from 0.291 to 8.26×10^-5^ with 16 variants having VAFs of >0.01, and 56 variants having VAFs of >0.001. All cultures also contained mtDNA insertions/deletions (indels) of 1 bp, >1 bp or more complex variants involving 2-19 bp that are readily identified by though not as efficiently captured or quantified by Duplex sequencing (**Figure 2C**). Among 25 mtDNA variants shared between at least two GSC cultures (16 SNV, 9 indel variants), a majority (18/25) were predicted not to alter coding and likely represent common population polymorphisms (**Tables S3, S4**).

GSC culture mtDNA SNVs were strongly biased to G>A/C>T transitions, with low levels of putative oxidative damage-linked G>T/C>A transversions: the few apparent oxidative damage-induced mutations may reflect in part continuous GSC growth in low (5%) oxygen. SNV variants were distributed over the whole mtDNA genome including the regulatory ‘D-loop’ region, with over half (59.2%) in coding genes where 67.4% were predicted to alter or terminate a protein open reading frame (**Figure 2D**). Subclones of mtDNA variants were identified in all cultures by ≥2 Duplex sequence reads containing the same mtDNA variant(s): these distinct molecular subclones represented ∼23% of 0827 reads, versus 8-10% of all reads in GBM4, GBM8 or 0131 (**Figure S3**). SNV molecular type distributions were similar across GSC cultures, and did not change appreciably after 1 - 2 Gy of IR (**Figure S4**). These results are the deepest look to date at mtDNA variation and subclonal architecture in GBM-derived GSC cultures, and resemble prior Duplex analyses of normal human brain mtDNA (Hoekstra et al., 2016; Kennedy et al., 2013; Williams et al., 2013).

### Gene expression profiling

GSC cultures GBM4 and GBM8 were the most similar in bulk and single-cell RNA sequencing analyses with 0131 was the most distinctive (**Figure 3A**), where UMAP plots of sparser scRNA data (**Figure 3B**) do a particularly effective job of capturing and displaying GSC-specific expression variance (Hsu and Culhane, 2023). These results are qualitatively similar to prior GBM gene expression data (Brennan et al., 2013; Wang et al., 2021, 2018a), and reported GSC-specific expression data for 0131, 0827 (Toledo et al., 2015) and for GBM8 (Wang et al., 2018b; Eyler et al., 2020). Flow cytometry and scRNAseq analyses documented a high growth fraction of putative cycling cells in all four GSC cultures (**Figures 1C, 3C, S5**).

**Figure 3.**
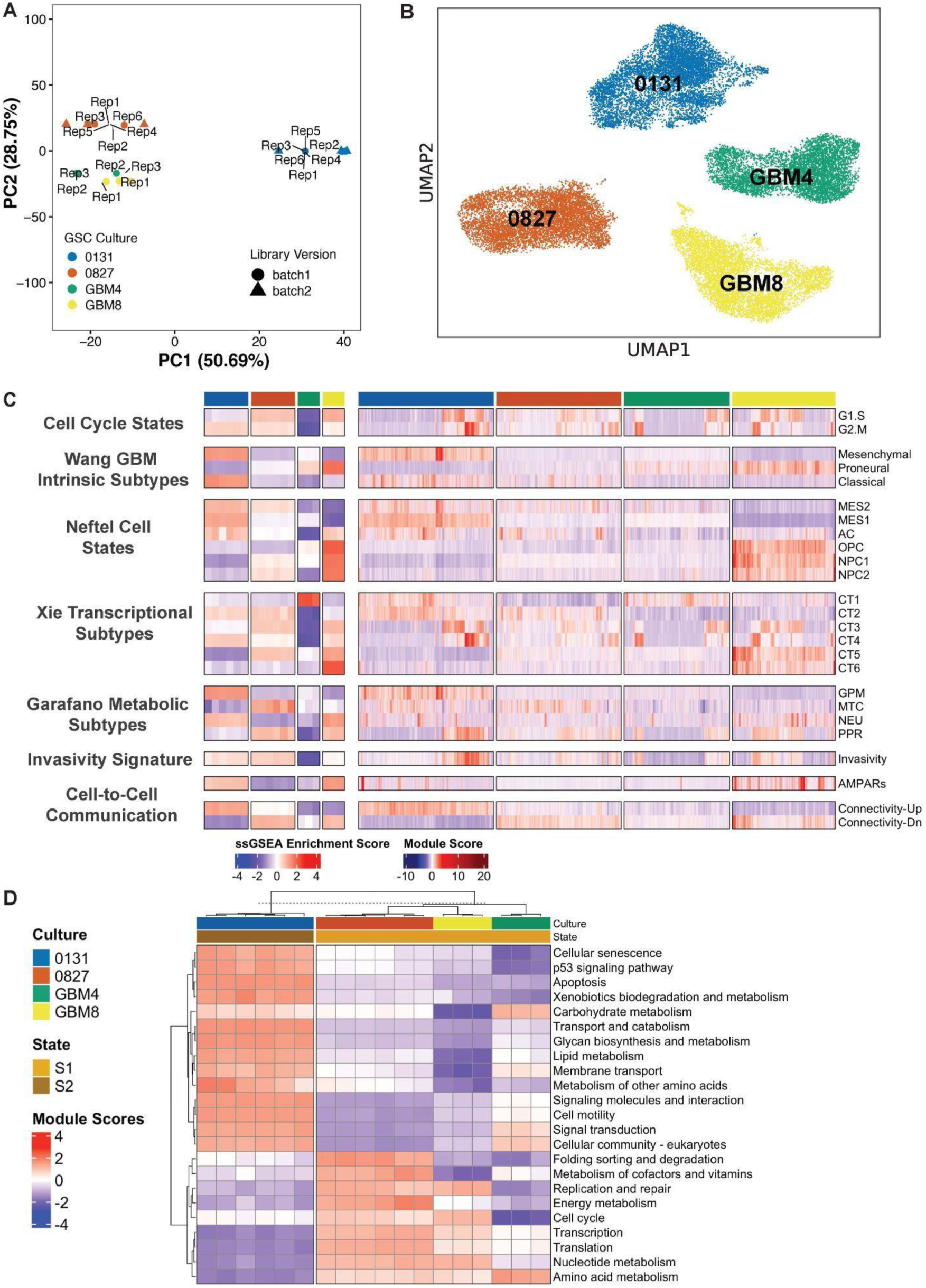
**Transcriptional profiling defines GSC cellular subtypes and states**. **(A)** PCA plot of gene expression in replicate bulk GSC samples, and **(B)** UMAP plot of scRNA-seq expression reveals transcriptome cellular heterogeneity. **(C)** Bulk (left) and single-cell (right) analysis data identify gene programs for cell cycling/growth fractions (as G1.S, G2.M phases), and quantifies expression of previously identified Wang GBM intrinsic and Garofano metabolic subtypes ((Wang et al., 2018a)(Garofano et al., 2021)) and cell states (Neftel ‘6’ and Xie CT1-CT6 states; (Neftel et al., 2019; Xie et al., 2024). Additional enrichment scoring for gene expression signatures reflecting potential/activity for invasivity, glutamate-mediated cell signaling via AMPAR iontropic transmembrane receptors and more for cell-cell connectivity and signaling. See text for additional detail and signature sources. **(D)** GSC culture gene expression data mapped onto a Functional Module States (FMS) framework reveals enrichment for metabolic pathways/processes including cell signaling, proliferation, metabolism, senescence and death. ***Abbreviation key:*** AC, astrocytic; GPM, glycolytic/plurimetabolic; MES1/2, mesenchymal states; MTC, mitochondrial; NEU, neuronal; NPC neural progenitor cell; OPC, oligodendrocyte progenitor cell; PPR, proliferative-progenitor related)(Garofano et al., 2021; Neftel et al., 2019; Xie et al., 2024)

Previously defined GBM transcriptional subtypes and cell states could be readily identified among our GSC cultures. For example, signatures developed by Neftel and colleagues (Neftel et al., 2019) identified strong enrichment for mesenchymal subtype expression states MES1/MES2 in 0131 and, to a lesser extent, an AC (astrocytic)-like state whereas GBM4, GBM8 and 0827 displayed proneural/NPC-like signatures (**Figures 3C, S6**). All four cultures expressed cell surface proteins and gene sets used to define GBM stem cells or stem-like behavior (e.g., *CD44, MYC, PROM1, CTNNB1, EZH2,* and *SOX2;* **Figures S5, S7, S8; Table S8**). Preliminary flow cytometry and fluorescence microscopy analyses documented heterogeneous cell surface marker expression: e.g., CD44 was strongly expressed by 0131 and 0827 and, at lower levels, in GBM4; with GBM8 expressing CD33/PROM1 but not CD44.

CD177/L1CAM expression was detected in GBM4, GBM8 and 0131, though not previously reported expression of CD15/FUT4 (**Figures S7, S8;** additional data not shown). The recently described transcriptional subtype/cell state hierarchy was also readily identified, from stem-like CT1 cells most abundant in GBM4, to unique, potentially post-mitotic CT-6 cells most abundant in GBM8 (Xie et al., 2024)(**Figure 3C**).

Proteo-metabolic and functional subtyping have also provided insight into GBM biology and therapeutic vulnerabilities (Oh et al., 2020)(Garofano et al., 2021; Kim et al., 2024; Migliozzi et al., 2023)(**Figure 3C**). GSC culture 0131 most strongly resembled the previously described glycolytic/plurimetabolic (GPM) GBM metabolic subtype, consistent with high activity in glycan biosynthesis (**Figure 3C, 3D**). In contrast, 0827 displayed mitochondrial/oxidative phosphorylation-dependent (MTC) features, consistent with high mitochondria-associated gene expression (**Figure 3C, S9**). GBM8 displayed both neuronal (NEU) and proliferative/progenitor (PPR)-like features, whereas GBM4 displayed greater MTC than PPR character (**Figure 3C**)(Garofano et al., 2021).

Gene expression programs also capture and can reflect important GBM non-cell-autonomous features (Cirigliano and Fine, 2025; Fine, 2024; Sloan et al., 2024). These include gene expression programs promote cell proliferation and invasion (Kim et al., 2024; Liu et al., 2024; Taylor et al., 2023; Venkataramani et al., 2019, 2022; Venkatesh et al., 2019); cell-cell communication and electrical coupling that may drive proliferation (Venkatesh et al., 2019)(Hausmann et al., 2023); and local as well as more distant tumor/non-tumor cell communication (Mangena et al., 2025; Venkataramani et al., 2022; Venkatesh et al., 2019; Zhang et al., 2025)(**Figure 3C**). GSC cultures, with the exception of GBM4, displayed expression and module enrichment scores favoring invasion (**Figure 3C**). Cell signaling and cell-cell communication/ synaptic coupling potential assessed by glutamate (or AMPAR) receptor expression was strongest in GBM8, followed by 0131 in connectivity ‘up/down’ signatures (Hai et al., 2024)**(Figure 3C**). Of note, GBM8 displayed the highest fraction of OPC-like cells, where elevated expression of both *PDGFAR* and *KCND2* expression have been linked to cortical bursting, synchronous electrical discharges and potential tumor-associated seizure activity (Zhang et al., 2025)

Functional Module States (FMStates) analysis (Qin et al., 2022) provided additional insight into GSC biology and behavior. Harmonized GSC and TCGA-GBM gene expression data were captured as three Functional Module factor-defined clusters with GBM4, GBM8 and 0827 displaying proneural, and 0131 mesenchymal, subtype features (**Figures 3D, S10**). GBM4, GBM8, and 0827 had high expression for amino acid metabolism, transcription and protein translation modules, consistent with high cell cycling potential. In contrast, 0131 displayed high expression of modules associated with cellular senescence, apoptosis, cell motility, cellular community, glycan biosynthesis and lipid metabolism and lower transcription/translation-associated module expression (**Figure 3D**). GBM4, GMB8 and 0827 displayed higher module scores for cell cycle, replication and repair, transcription, translation, and amino acid metabolism, and lower scores for community and amino acid metabolic activities (**Figure S10**).

### Proteomic and phosphoproteomic response to radiation

MS-MS proteomic profiling of GSC cultures identified ∼4,000 protein groups and ∼10,000 phosphopeptides both prior to and after IR treatment with high (R >0.9) reproducibility (**Figure 4, S11 and S12**). In order to assess the proteomic response to IR, we treated GSC cultures with 1 Gy of IR followed by sampling at 1 hr to capture the early-intermediate IR response: pilot data indicated 1 Gy of IR strongly induced γ-H2AX, a DNA damage response indicator, at 1 hr while leaving a majority of cells still viable 24 hrs later (**Figure 1E**).

**Figure 4:**
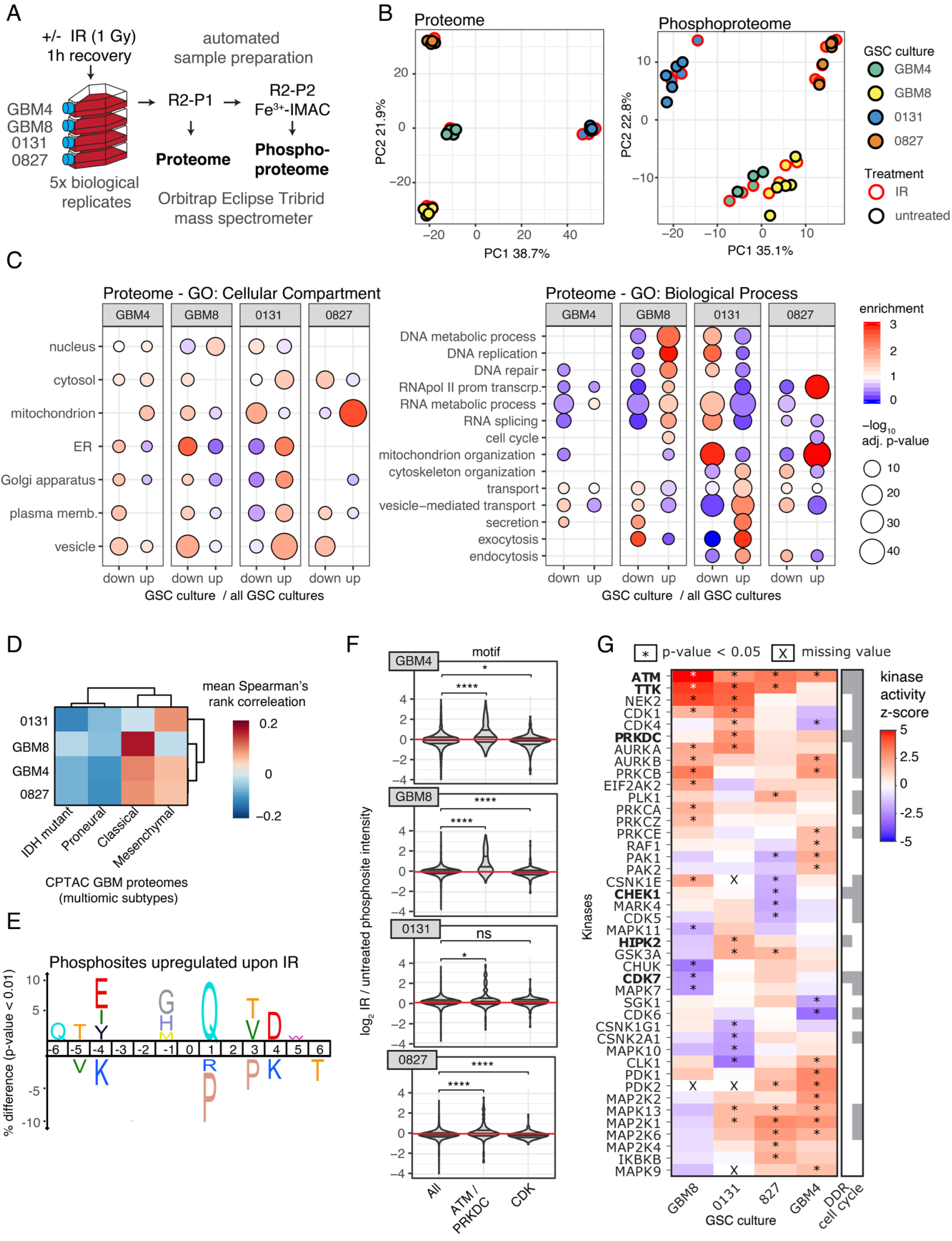
GSC culture proteomic/phosphoproteomic profiling by mass spectrometry. (**A**) Workflow for mass spectrometry-based profiling using rapid-robotic (phospho)proteomic sample prep protocols (R2-P1/P2)(Leutert et al., 2019). (**B**) Comparative PCA plots of GSC-specific protein/phosphosite abundances as a function of IR. (**C**) Gene Ontology (GO) term enrichment analysis of GSC-specific changes by cellular compartment (left) and biological process (right). (**D**) Mean Spearman rank correlations between GSC proteomes and untreated GBM samples grouped by IDH and Wang proteogenomic subtypes (Wang et al., 2021). (**E**) Amino acid enrichment/ depletion surrounding protein phosphosites in IR-treated GSC cultures. (**F**) Violin plots of log_2_-fold changes of phosphosite abundance in control versus IR-treated cells. All phosphosites (left column), and phosphosites corresponding to ATM/PRKDC (center) or CDK (right column) consensus motifs, where Wilcoxon tests of group median comparisons identify 0131 as an outlier. (**G**) Kinase substrate enrichment analysis (KSEA) identifies IR-induced phosphosite changes. Negative/ positive z-scores imply predicted decrease/increase, respectively, in kinase overall activity in IR-treated versus control cells. Significant differences, corrected for multiple testing, are noted by asterisks (*). Gray scale code (shown right) identifies kinases associated with DNA damage response (DDR), cell cycle regulation (cell cycle), or both pathways.

Principal component analysis (PCA) of protein and phosphosite abundances displayed tight clustering in biological replicates, with clear separation among GSC cultures (**Figure 4B**).

GBM8 and 0131 most closely resembled classical and mesenchymal proteogenomic GBM subtypes, respectively, with GBM4 and 0827 displaying features of both classical and mesenchymal subtypes (Wang et al., 2021)(**Figure 4D**). All four cultures expressed both IDH1 and IDH2 proteins with the highest steady state IDH peptide levels in GBM8 and 0131. GBM8 also had the highest content of nuclear proteins associated with Gene Ontology (GO) DNA and RNA metabolic processes, a pattern also observed but less prominent in GBM4 (**Figure 4C**).

The highest content of cytosolic, ER and membrane proteins associated with secretory processes was found in 0131; 0827, in contrast, showed the strongest enrichment for mitochondrial and transcription-associated proteins (**Figure 4C**). Significantly altered phosphoprotein abundances were identified between 0827 and 0131 by unsupervised hierarchical clustering of untreated cultures (**Figure 4B**), and significant differences in the abundance of protein complexes associated with chromatin remodeling, DNA replication, cell cycle control, DNA mismatch repair and POL3-dependent transcription (**Figure 4C**).

Proteo-metabolic classes that are predictive of GBM therapeutic response could be identified in GSC proteomic data (Oh et al., 2020) (Garofano et al., 2021; Migliozzi et al., 2023): untreated 0827 cultures displayed different relative mitochondrial protein abundances, versus the elevated levels of oxidative phosphorylation-related proteins in GBM4 (**Figure 4C**). The most IR-sensitive cultures, 0827 followed by GBM4, displayed elevated expression of oxidative phosphorylation-related proteins. Reduced expression of the poor prognostic biomarker FKBP prolyl isomerase 9 (FKBP9) was observed in 0827 and, to a lesser extent, GBM4 with no corresponding trend for the favorable prognostic biomarker phosphoglycerate dehydrogenase (PHGDH).

IR treatment led to numerous protein phosphorylation changes, most notable among DNA damage response proteins and phosphorylation response pathways (**Figure S12**). These changes included significant post-IR [pS/pT]Q motif enrichment (**Figure 4E)** with quantitative induction in all cultures except 0131 (**Figure 4F**): the [pS/pT]Q and [pS/pT]Px[R/K] motifs correspond respectively to DNA damage-responsive ATM and/or DNA-PK consensus motifs, and to CDK kinase consensus motifs (Johnson et al., 2023)(**Figure S12C**). Many other IR-modulated phosphosites linked to specific biological functions were also identified (**Figure S12D**). For example, IR-induced ATM S1918 autophosphorylation both activates and targets the ATM kinase to DNA damage sites (Berkovich et al., 2007). GBM4 and GBM8 also displayed many additional IR-induced DNA damage response protein phosphorylation site changes, e.g., MDC1 phosphorylation, a regulator of the DNA damage response, was observed in all cultures except 0131. IR-induced phosphorylation site changes in XRCC4, RAD9 and H2AFX, in contrast, were observed only in 0131, our most radio-resistant GSC culture.

Kinase-Substrate (KSEA) Enrichment Analysis (Wiredja et al., 2017) identified additional IR-regulated phosphorylation site-kinase relationships. ATM was among the top predicted active kinases across all GSC cultures; was most active in GBM8; and was followed by TTK in all but GBM4, or by NEK2 and CDK1 in GBM8 and 0131 (**Figure 4G**). Several activated MAPK members were predicted in GBM4 and 0827, as were differential IR activation of protein kinase C isozymes, DNA-PKcs and PAK1/PAK2.

All of these kinases have recently been linked to functional GBM proteo-genomic subtypes, with DNAPK/PKRDC as an apical or ‘master’ kinase regulating many GBM phenotypic hallmarks (Migliozzi et al., 2023). DNAPK/PKRDC acts together with PTPN11 in a protein tyrosine phosphatase regulatory signaling hub in high grade gliomas (Liu et al., 2024). We identified the key PTPN11 regulatory phosphosite, Y62, and significant post-IR phosphorylation changes in all four GSC cultures, with 0827 having the highest Y62 phosphorylation levels associated with an activating *EGFR* mutation (Toledo et al., 2015). PTPN11-S591, a phosphosite with few known functional roles, was also identified though not PTPN11-Y546 which has been reported in *IDH*-mutant gliomas (Liu et al., 2024). These proteomic, genomic and gene expression data collectively characterize GSC-specific features that should allow more informed selection among cultures for experimental use.

### GSC population heterogeneity and post-IR cellular trajectories

We used mtDNA variants together with expressed lentiviral DNA barcodes (also known as ‘CellTags’, (Biddy et al., 2018)) and single cell RNA sequencing to characterize and track GSC cellular heterogeneity over 40 days after IR treatment. Our experimental design (**Figure 5A**) used IR doses predicted to kill ∼80% of GSC cells, where experimentally determined cell killing ranged from 83% to 65% 21 days after IR treatment. We reasoned this level of cell killing might allow the identification of initial gene expression signatures linked to cell loss or survival, while avoiding the dominant, potentially less informative ‘jackpot’ subclones that often follow high fractional cell kills and/or repeated treatment cycles (see, e.g., (Eyler et al., 2020)).

**Figure 5:**
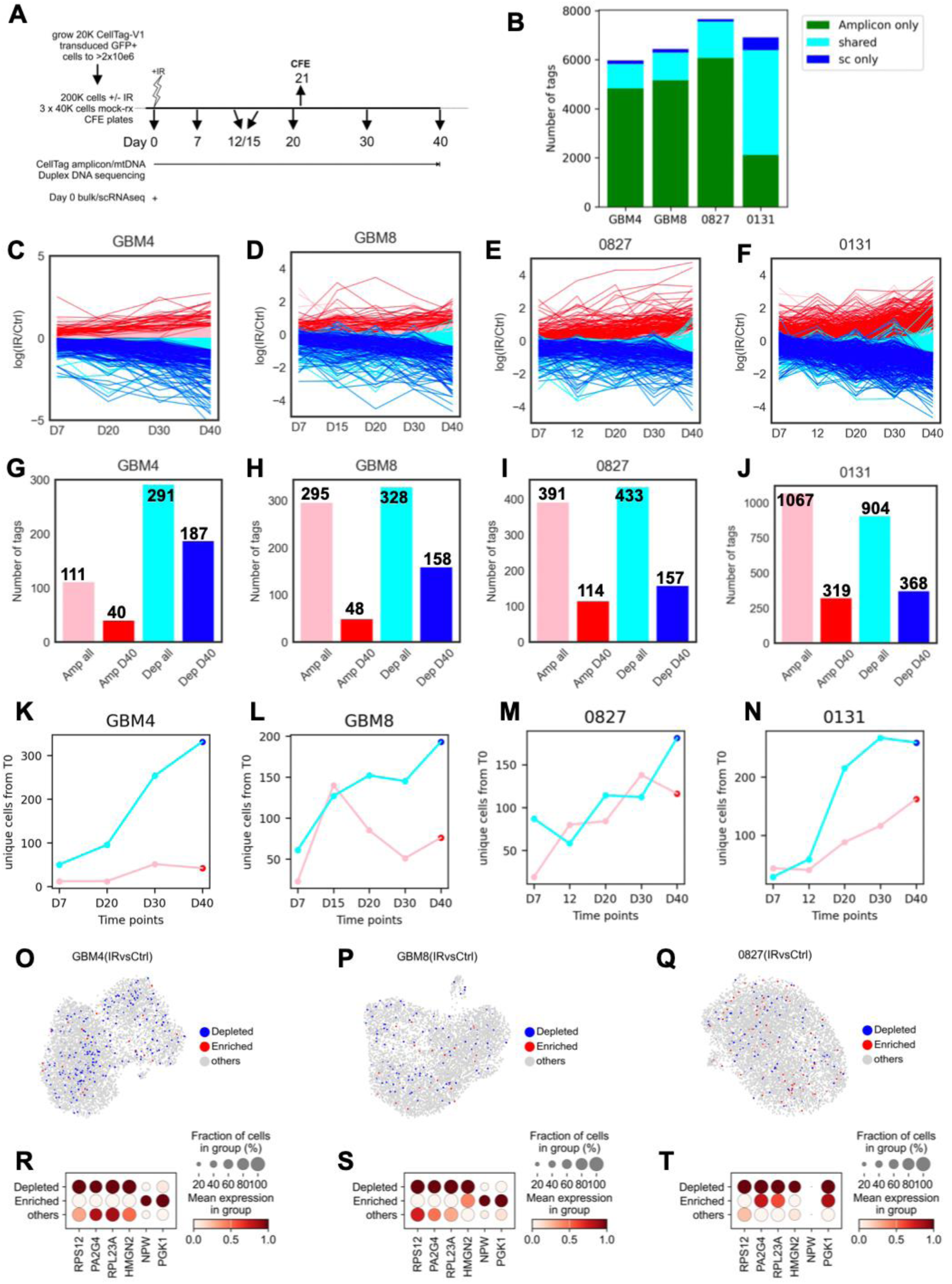
Molecular barcode identification of cellular heterogeneity and trajectories after IR treatment. (**A**) Design of 40 day time course sampling experiment to follow lentiviral CellTag and mtDNA variant frequencies in control (-IR), irradiated (+IR) and mock-treated, reduced complexity (mock) experimental arms (see Methods for additional details). (**B**) CellTags identified in GSC cultures by amplicon sequencing (green), scRNA sequencing (blue) or both sequencing methods (teal). (**C-F**) Unique sequence CellTag trajectories over 40 days ± IR identify subsets with significant enrichment (red), potential enrichment (pink), depletion (blue), potential depletion (cyan) or no significant change (gray) by Wilcoxon test p-value of IR/control ratios. (**G-J**) Unique sequence enriched/depleted CellTags across multiple time points (all), or Day 40 vs. Day 0 (D40). (**K-N**) Unique cells with at least half of cell-associated CellTags enriched (*pink*) or depleted (*cyan*) by at least 50% at D40. (**O-Q**) Cells enriched (red) or depleted (blue) as a function of IR treatment mapped onto all cells identified by scRNAseq. (**R-T**) Fraction of cells and level of expression of genes differentially expressed in enriched vs. depleted cells as a function of IR treatment. At least two of three GSC cultures were required to display the same associations to be included. A comparable analysis of differentially expressed genes in enriched versus depleted cells in control cultures is shown in **Figure S16**.

GSC culture heterogeneity was assessed by initial bulk and single cell gene expression analyses coupled with mtDNA variant and lentiviral molecular barcode profiling (‘CellTags’, after (Biddy et al., 2018), hereafter termed ‘tags’)(**Figure 5A, 5B**). Both tag and mtDNA sequence variants were then tracked across multiple sampling timepoints and linked back to individual cells identified by their UMIs in scRNAseq data. This strategy allowed us to track individual cells and their descendants over time, and to link starting gene expression programs to individual cellular trajectories (**Figure S14, S15**). Lentiviral tagging generated ∼ ⅓ of cells in three GSC cultures with mean tag counts of 2 to 3/cell (range 2.17 - 3.34), with 0131 an outlier with over 90% of cells tagged and a mean tag count of 17.35. Mean unique tags linked to a cell-identifying UMI ranged from 1.33-6.04 (**Figure S13, S15**).

Tag counts allowed us to identify five trajectory classes in untreated GSC cultures by K-means clustering using corrected, log-transformed ratios of time point/Day 0 counts (**Figure S16A-D**). IR-enriched or-depleted cells containing these tags were then identified in IR-treated cultures by a Wilcoxon test for p-values of P<0.1 and tag read ratios of IR/control of >2 (for enriched tags) or <0.5 (for depleted tags)(**Figure 5C-F, Figure S16E-G**). Tag and cell depletions were more common than enrichment, most notably in GBM4 and GBM8 (**Figure 5G-N**). Gene signatures shared across at least two cultures identified 16 genes associated with tag enrichment or depletion in untreated cells, and 6 in IR-treated cultures (**Figure 5R-T, S16H-J**). Top genes identified in GBM4 and GBM8 independent of IR treatment were involved in ribosomal or mitochondrial structure and function (**Figure 5R-T, Figure S16H-J**), whereas in IR-treated cells the top genes regulated gene structure/expression or ribosome structure (e.g., RPL23A and RPS12)(**Figure 5R-T**): the up-regulation of ribosome-related genes suggests that cells with higher inferred ribosomal gene expression may be more IR-sensitive. A contrast is provided by mitochondria-related genes such as MT-CO3, MT-ND3, MT-ND4, MT-CYB that were up-regulated in enriched cells, suggesting a role for mitochondrial activity in IR-treated survival and proliferation. We attempted but could not perform a comparable analysis using expressed mtDNA variants, in light of their low frequency in GBM4, GBM8 and 0131 cultures (range 0.13-0.19% of cells)(**Figure S4, S17**).

## Discussion

We have integrated new genomic, proteomic and phenotypic data on four IDH+ glioma stem cell (GSC) cultures to serve as tractable cellular models of the most common genomic subtype of adult primary glioblastoma (GBM). The four GSC cultures from unrelated adult donors differed in genomic, transcriptomic, proteomic and therapeutic response features, though collectively reflected previously reported subtypes and features of primary GBM neoplastic cell populations (Lasorella and Iavarone, 2021; Taylor and Monje, 2022). These results should make it easier to identify the best GSC culture(s) to use to investigate specific questions in GBM biology, including the mechanisms driving persistence, therapeutic resistance and recurrence (Kim et al., 2024; Liu et al., 2024).

All four GSC cultures that we characterized were initiated from GBM tumor resection tissue that was histologically classified as glioblastoma (Son et al., 2009; Wakimoto et al., 2009). Each was markedly aneuploid with regional and gene-specific amplification or deletion (e.g., *MYC* amplification in GBM4, and *MYCN*, *PDGFRA* and *MDM2* amplification with homozygous *CDKN2A/B* deletion in GBM8). Whole exome sequencing (WES) identified 130 variants in genes linked to GBM risk or pathogenesis, with from 3 variants in 2 genes (GBM8) to 36 variants in 17 genes (0827) in the top 25 most frequently altered genes in GBM. The hundreds of GSC mtDNA variants we identified by sensitive, high accuracy Duplex DNA sequencing represent the deepest look to date at mtDNA variation in GBM and in GSC cultures. Duplex DNA sequencing provides a more rapid way to identify, count and determine the subclonal architecture of these cell-intrinsic molecular barcodes and their use, e.g., for lineage tracking, than alternative approaches such as cellular subcloning (e.g., (An et al., 2024)). Duplex DNA sequencing of mtDNA variants thus powerfully augments a useful and revealing general strategy also being explored by others (Kim, 2023; Ludwig et al., 2019; Nitsch et al., 2024; Sankaran et al., 2022).

Gene expression profiling confirmed the close resemblance of our GSC cultures to previously defined GBM transcriptional subtypes: 0131 displayed strong enrichment for mesenchymal states MES1/MES2, whereas GBM4, GBM8 and 0827 displayed proneural/NPC-like states (Neftel et al., 2019). All four cultures variably expressed cell surface proteins and gene sets that identify or define GBM stem cells or stem-like behavior (e.g., *CD44, MYC, PROM1, CTNNB1, EZH2,* and *SOX2).* Transcriptional subtypes that suggest a cell state hierarchy were also identified, from stem-like CT1 cells that were most abundant in GBM4, to unique potentially post-mitotic CT-6 cells that were most abundant in GBM8 (Xie et al., 2024). Among GBM metabolic subtypes GSC 0131 most strongly resembled the glycolytic/plurimetabolic (GPM) metabolic subtype, whereas 0827 displayed mitochondrial/ oxidative phosphorylation-dependent (MTC) features. GBM8 had both neuronal (NEU) and proliferative/progenitor (PPR)-like features, with GBM4 displaying greater MTC than PPR character (Oh et al., 2020)(Garofano et al., 2021; Kim et al., 2024; Migliozzi et al., 2023).

Parallel MS-MS analyses provided deep proteomic data that complemented and extended these genomic and gene expression changes in steady-state and IR-treated cultures. ATM was the top predicted active kinase across all GSC cultures in Kinase-Substrate (KSEA) Enrichment Analysis (Wiredja et al., 2017), followed by TTK in all but GBM4, and NEK2 and CDK1 in GBM8 and 0131. MAPK members were active in GBM4 and 0827, with post-IR activation of protein kinase C isozymes, DNA-PKcs and PAK1/PAK2. We also identified DNAPK/PKRDC, recently identified as a ‘master’ kinase and part of a regulatory signaling hub with the tyrosine phosphatase PTPN11 in high grade gliomas (Liu et al., 2024). All of these kinases have been linked to functional GBM subtypes (Migliozzi et al., 2023).

GSC cultures hold particular promise in the search for new GBM therapies and treatment regimens. All four of our GSC cultures displayed highly reproducible dose-survival curves for the GBM standard-of care therapies of IR and temozolomide (TMZ). IR sensitivity could be reproducibly enhanced by pre-treatment with ATM kinase inhibitors, though TMZ sensitivity did not consistently reflect MGMT protein expression as others have noted (see, e.g., (Nagel et al., 2017). We further investigated these discrepancies by growing GSC cultures in the demethylating base analog decitabine (5-aza-2′-deoxy-cytidine) to induce MGMT expression (Moen et al., 2014). However, we found that decitabine killed all four GSC cultures at sub-micromolar concentrations, a potent cytotoxic activity that may be therapeutically useful in conjunction with, e.g., GBM-targeting immune therapies: see, e.g., (Silva-Hurtado et al., 2024; Eramo et al., 2005; Ma et al., 2022; Patties et al., 2019). We also found that the microtubule-binding drug ST-401, already shown to be active against GBM xenografts, killed GSC cultures at low nM concentrations (Horne et al., 2021). Of note, this small molecule acts by a different mechanism of action than colchicine or nocodazole (Vicente et al., 2024). These examples indicate how GSC cultures can be used to rapidly establish potency, identify potential mechanisms of action, and search for useful synergies with other drug candidates (e.g., vortioxetine, (Lee et al., 2024) and gliocidin (Chen et al., 2024)) or IR to be tested in more complex GBM disease models.

Several general features of GSC cultures need to be kept in mind when contemplating experimental use. First, early passage GSC cultures often consist of mixtures of neoplastic and non-neoplastic cell types, where there is rarely a patient-specific, non-neoplastic control sample from the same patient. This can in part be addressed by using unrelated, non-neoplastic neural stem cell cultures as controls in conjunction with experimental designs that compare treated versus untreated cultures (Toledo et al., 2015). GSC cultures are variably aneuploid and likely genomically unstable, though there are few datasets to help quantify instability at different genomic levels. Despite this, many GSC chromosome-level and smaller genomic changes appear to be relatively stable in culture and may collectively support continuous proliferation (Toledo et al., 2015; Wakimoto et al., 2012). Many of these and related experimental issues can be minimized by using replicate, minimally passaged GSC cultures for experimental series, supplemented by single cell analysis methods and readily generated, clonally derived subcultures.

The growing use of GSC cultures as cellular GBM disease models reflects their unambiguous origin from human disease tissue, experimental tractability and versatility when compared with other GBM disease models. GSC cultures are especially useful for rapid, high throughput profiling of ‘therapeutic space’ (see, e.g., (Bashi et al., 2024)) versus large libraries of chemicals or gene targeting agents. These screens can be further sensitized and targeted by using engineered isogenic GSC gene-and allelic series. The same GSC cultures can then be used to initiate a growing array of ‘on-chip’, xenograft and organoid protocols to allow analyses of both conventional and newer therapies including antibodies, antibody-drug conjugates (ADCs) and cellular therapies: see, e.g., (Dundar et al., 2020; Furnari et al., 2024; Linkous et al., 2019)(Akter et al., 2021; Robertson et al., 2019; Sahu et al., 2022). Perhaps surprisingly, GSC cultures can also be used to investigate some tissue-level, non-cell-autonomous features of GBM (Huang-Hobbs et al., 2023; Mangena et al., 2025; Sloan et al., 2024; Taylor and Monje, 2022; Varn et al., 2022; Venkataramani et al., 2022). For example, the compact, nodular growth of GBM4 mouse orthotopic xenografts is predicted from their low invasivity score, versus the predicted diffusely invasive xenograft growth of GBM8 (Wakimoto et al., 2009).

All of these examples highlight how GSC culture experimental tractability and versatility are leading to imaginative new uses to advance GBM biology and translational science while addressing both general and GSC-specific weaknesses as cellular disease models (Ghandi et al., 2019; Gillet et al., 2013; Gutmann et al., 2025). Thus the continued development and imaginative use of GSC cultures should improve our understanding of GBM biology, and aid the rapid identification of more effective GBM therapies and treatment regimens (Aldape et al., 2019; Fine, 2024)(Sloan et al., 2024).

## Acknowledgments

We thank the following individuals for help, reagents and discussion during the course of these experiments: Andres Barria, Amy Hong, Theo Knijnenburg, Andrei Mikheev, Colin Pritchard, Robert Rostomily, Nephi Stella, Tom Walsh and Piri Welsch. Christine Disteche interpreted GSC culture karyotypes. Jeongwu Lee (Cleveland Clinic Center for Cancer Stem Cell Research) and P.J. Cimino (Neuropathology Head, NIH Surgical Neurology Branch) provided additional information allowing us to more clearly understand the origins of GSC culture 0827. We thank Brian Whitmire and Melissa Randell at 10X Genomics for facilitating access to their single cell sequencing reagents, and Jose McFaline-Figueroa and Dana Jackson of the Trapnell Lab (UW Genome Sciences) for expert guidance in implementing scRNAseq analyses. Erica Jonlin (UW Institute for Stem Cell and Regenerative Medicine) provided expert counsel on the appropriate experimental use of these patient-derived, anonymized GSC cultures.

## Authorship

(14 Contributor Roles Taxonomy (CRediT: docs.casrai.org/CRediT) categories are combined here under 6 broad headings):

## Conceptualization

NC, SRK, ML, RM, GQ, IS, BT

## Funding/Resources

JG, SRK, RM, IS, JV

## Investigation (Data curation, Formal Analysis, Methodology, Software, Validation, Visualization)

NC, SRK, BK, ML, RM, GQ, MS-C, IS, WT, BT

## Project administration/Supervision

SRK, RM, IS

## Writing: original draft

NC, SRK, ML, RM, GQ, BT

## Writing: review & editing

NC, JG, SRK, BK, PL, ML, RM, PP, GQ, MS-C, WT, BT, JV

## Dedication

We dedicate this manuscript to our late colleague and co-author Ilya Shmulevich. Ilya played a key role in initiating and shepherding this project, and in advancing so many areas of cancer genomics as collaborative, open science with publicly accessible resources. We sorely miss his brilliance, insight, humanity and humor.

## DNA/RNA sequencing data

These data and records, archived under NCBI SRA Bioproject PRJNA1036631 (https://www.ncbi.nlm.nih.gov/sra/PRJNA1036631), will be available upon publication: whole exome sequencing data, SRA SUB15419289; Duplex mtDNA sequencing data, SRX24964902-SXR24964933; bulk RNAseq data, SRA SUB15367655 (raw reads) and GEO GSE303069 (processed data); single-cell RNA-seq data, GEO GSE303662; and CellTag molecular barcode amplicon sequencing data, GEO GSE302554.

## MS-MS proteomic data

The mass spectrometry proteomics and phosphoproteomics data have been deposited with dataset identifier PXD035886 to the ProteomeXchange Consortium via the PRIDE Partner Repository (Citation: https://doi.org/10.1093/nar/gkab1038).

## Code availability

In-house developed bioinformatics pipelines and code can be found at the following Github repositories: For version 2.1.2 of whole exome and mtDNA Duplex data processing pipelines and code, see: https://github.com/Kennedy-Lab-UW/Duplex-Seq-Pipeline For gene expression and amplicon bar code analysis pipelines and code, see: https://github.com/IlyaLab/GSC_culture_characterization

