## Supplement for "Cellular heterogeneity and therapeutic response profiling of human IDH+ glioma stem cell cultures"

**New Results**

[a] equal contributing co-authors listed alphabetically, with respective contributions of all authors detailed in the CRediT Taxonomy found after Acknowledgements.
[1] Institute for Systems Biology, Seattle WA
[2] Institute of Molecular Medicine, Feinstein Institutes for Medical Research, Manhasset NY
[3] Department of Molecular Medicine, Zucker School of Medicine at Hofstra/Northwell, Hempstead NY
[4] University of Washington, Departments of Laboratory Medicine and Pathology (SK, MS-C and RM); Genome Sciences (ML, RM, JV); Bioengineering (RM) and Neurology and Neurosurgery (JG), Seattle WA
[5] Fred Hutchinson Cancer Research Center, Seattle, WA
*current address: Qualitel, Everett WA (PL); Roche Pharma R&D, Basel, Switzerland (ML); and Umoja Biopharma, Seattle WA (WT)
[≈] these authors contributed equally to this manuscript
[†] deceased

**Keywords/indexing terms:** glioblastoma, glioma stem cell, ionizing radiation, temozolomide, whole exome sequencing, single cell RNA sequencing, MS-MS proteomic profiling, cellular heterogeneity, mtDNA variant, Duplex DNA sequencing, expressed lentiviral DNA barcode.

**Creative Commons Attribution: CC BY-NC 4.0**

**Supplementary Materials - Index**


1. **Extended Methods**

2. **Supplementary Figures**

   **Figure S1: GSC dose-response profiles for ionizing radiation, temozolomide and additional small molecules.**

   **Figure S2. GSC cultures are aneuploid with numerous numerical and structural chromosome abnormalities.**

   **Figure S3: GSC mtDNA mutational spectrum and subclonal structure identified by Duplex DNA sequencing.**

   **Figure S4: GSC mtDNA DNA base substitution variant (SNV) frequency, spectrum and subclonal composition as a function of time and IR treatment.**

   **Figure S5. GSC culture cell cycling metrics.**

   **Figure S6: GSC cultures exhibit transcriptionally-defined GBM cellular states.**

   **Figure S7: GSC culture expression of markers of stemness.**

   **Figure S8: Expression of GSC surface markers and key functional biomarkers.**

   **Figure S9: GSC 10X scRNA sequencing data prior to and after filtering.**

   **Figure S10. Comparison of GSC and TCGA-GBM samples using Functional Module states scores based on gene expression.**

   **Figure S11: Additional details on GSC proteomic and phosphoproteomic characterizations including DNA metabolic protein complexes.**

   **Figure S12: GSC proteomic and phosphoproteomic response to ionizing radiation.**

   **Figure S13: CellTag labeling and distribution in GSC cultures.**

   **Figure S14: Total and unique CellTag barcode reads across GSC cultures, experimental arms and sampling time points.**

   **Figure S15: Single cell identification and relatedness identified from expressed, unique-sequence CellTag barcodes.**

**Figure S16: Molecular barcode trajectories in untreated GSC cultures reveal tag and cellular enrichment/depletion over 40 days in cultures.**

**Figure S17: mtDNA variant trajectories over 40 days in control and IR-treated GSC cultures.**

#### 3. Supplementary Tables

**Table S1: Short tandem DNA repeat allele length signatures unambiguously identify GSC cultures.**

**Table S2: Single nucleotide (SNV) variants identified by GSC culture exome sequencing.**

**Table S3: GSC culture Duplex mtDNA sequencing metrics.**

**Table S4: GSC culture-specific mtDNA variants identified by Duplex mtDNA sequencing.**

**Table S5: Cell Ranger summary statistics from GSC culture scRNAseq analyses.**

**Table S6: Distribution of gene expression-defined GBM subtypes from single cell gene expression profiling data.**

**Table S7: Expressed mtDNA variants and expressed CellTags identified in GSC cells by scRNAseq UMI indexing.**

**Table S8: Gene sets for transcription functional characterizations.**

**Table S9: CellTag reads for time course sampling experiment.**

**Table S10: Gene set enrichment statistics for bulk and scRNAseq data analyses.**

**Table S11: Protein and phosphoprotein abundances and statistics**

### 1. Extended Methods

#### Glioma stem cell cultures

GBM4 and GBM8 were initiated from surgical resection tissue from unrelated adult GBM patients as previously described in (Wakimoto et al., 2009), or for 0131 and 0827 as previously described in (Son et al., 2009). GSC cultures GBM4 and GBM8 cultures were provided by Drs. Robert Rostomily and Andrei Mikheev (University of Washington and Houston Methodist Hospital (GBM4 and GBM8), and 0131 and 0827 by Dr. Patrick Paddison (Fred Hutchinson Cancer Center, Seattle WA). All four cultures have alternate designations as published, and/or in the Cellosaurus database: GBM4/MGG4 = RRID:CVCL_D1H1; GBM8/MGG8 = RRID:CVCL_D1H4; 0131 = RRID:CVCL_C9EE; and 0827 = RRID:CVCL_C9EF. We use for consistency and brevity the designations GBM4, GBM8, 0131 and 0827 throughout.

GSC 0131 was from an untreated adult female, whereas 0827 was from a recurrent GBM as revealed by analyses here that identified a high tumor mutational burden (TMB) and prominent TMZ-associated single base mutational signature (see Results). These findings, together with additional clinical and pathologic data provided by Drs. Jeongwu Lee (Cleveland Clinic Center for Cancer Stem Cell Research) and P.J. Cimino (Neuropathology Head, NIH Surgical Neurology Branch) helped confirm the identity of 0827 as a recurrent, previously treated GBM-derived GSC culture.

GSC culture sex assignments were confirmed by a combination of karyotyping, comparative genomic hybridization and/or molecular fingerprinting (Wakimoto et al., 2009; Wang et al., 2018), but could not be independently confirmed in light of IRB restrictions preventing the disclosure of patient ages and sexes (Drs. Dr. Haroaki Wakimoto and Howard Fine, personal communications). In light of the absence of demographic or patient-specific data beyond what has been published, and the use of anonymized human tumor cell cultures derived from deceased individuals, this work is not considered human subjects research as defined by federal and state guidelines.

New short tandem repeat (STR) DNA authentication profiles were established for all four cultures, as none were found in the DSMZCellDive (Koblitz et al., 2022) or CLIMA 2.2 (Romano et al., 2009) STR authentication databases (both re-accessed 6 March 2025). STR profiles were generated using the CellCheck 9 Plus STR panel (IDEXX BioAnalytics, Columbia, MO) that includes additional primers to exclude common cell line and species contaminants. Cultures were repeatedly PCR-screened, and shown to be negative for cell line, species and *Mycoplasma* spp. infection.

GSC cultures were propagated in serum-free medium at 37°C in a low oxygen (5% $O_2$/5% $CO_2$) humidified atmosphere on untreated cell culture plastic. Single cell suspensions were generated using Accutase™ with pipetting to generate single cell suspensions for hemocytometer counting and dilution (typically 1:10 or 10-fold) into complete growth media and new flasks. GBM4 and GBM8 were grown in Neurobasal medium (ThermoFisher) supplemented with B-27 and N-2 according to manufacturer instructions (ThermoFisher), EGF (PeproTech, 20 ng/mL), FGF (PeproTech, 20 ng/mL) and heparin (Sigma, 5 ug/mL). GSC cultures 0131 and 0827 were grown in Neurocult medium (StemCell Technologies, Inc) supplemented with recombinant human EGF (20 ng/mL), FGF (20 ng/mL) and heparin (0.8 ug/mL). Large initial expansions were generated and used for STR-authentication, *Mycoplasma* testing and the establishment of archival replicate nitrogen stocks and working cultures.

**GSC culture proliferation rates and colony forming efficiencies**

Cell proliferation rates were determined by serial cell counting over 6 days after replating as described (Gui et al., 2016). Colony-forming efficiencies (CFE) were determined by dilution cloning in replicate 96-well plates, where 2 - 256 cells/well in 200 ul of complete growth medium were seeded into the central 60 wells, then surrounded with sterile water-filled wells to minimize edge evaporation artifacts. Plates were grown without refeeding for 2 - 3 weeks, then visually scored to enumerate empty wells and wells containing colonies of ≥50 cells. Colony forming efficiency was calculated from colony-negative and -positive well counts conditioned on the cell seeding density using the following formula: $CFE = -\frac{ln(P_o)}{n}$, where $P_o$= the decimal fraction of 60 wells that did not contain colonies (negative wells), and n = the average number of cells plated/well). Cell cycle phase distributions were determined by flow cytometric analysis of fixed and DAPI-stained exponentially growing GSC cultures as previously described (Dhillon et al., 2007).

**Metaphase Cytogenetics**

G-banded metaphase karyotypes were generated by the University of Washington Cytogenetics and Genomics Laboratory Research Service using a standard protocol to generate metaphase-arrested cells (Schreck and Distèche, 2001). All karyotypes were reviewed by an ABMG-certified cytogeneticist with extensive experience analyzing human clinical and cell line data. A minimum of 20 cells/culture were completely analyzed.

**Cell proliferation and survival as a function of IR or drug/small molecule dose**

IR suppression of GSC proliferation and survival was assessed using both bulk population suppression and colony-forming efficiency (CFE) assays. Bulk population assays used 500 - 1000 cells/well seeded in 200 ul of complete growth medium in 96 well plates as described above, then incubated for 3 hrs in the presence/absence of inhibitor (if used) in open Zip-lock™ bags inside a 5% $CO_2$/5% $O_2$ incubator. Sealed bags were irradiated in a RS-2000 X-irradiation unit (Rad Source Technologies, Buford, GA) at a dose rate of ~180 cGy/min to achieve a final dose range from 0 to 6 Gy. Irradiated and control plates were grown for 7-10 days after IR without refeeding, followed by WST-1 straining to quantify relative cell numbers in IR-treated and control wells. CFE assays were performed in duplicate 96 well plates with 2 - 256 cells/well seeded in 200 ul of complete growth medium as described above. ATM kinase inhibitors KU-60019 or KU-59933 were tested to determine their ability to modify IR dose-dependent population growth or CFE by inhibitor addition 3 hrs prior to irradiation, followed by 7-10 days or 2 - 3 weeks of undisturbed growth prior to quantifying cell number or counting wells containing colonies with ≥ 50 cells.

The ability of temozolomide to suppress GSC proliferation was determined using bulk population and CFE protocols as above, in conjunction with Western blot analysis and gene expression profiling to determine methylguanine methyltransferase (MGMT) expression. MGMT protein was detected in Western blots performed as previously described (Swanson et al., 2004) using a CellSignaling MGMT Rabbit 2739 primary antibody and a CellSignaling Anti-Rabbit IgG (7074) secondary antibody. The small molecules 5-aza-2'-deoxycytidine (decitabine); ST-401 and glutamate were analyzed to determine their ability to modify GSC proliferation and CFE as described above. In pilot experiments we also determined the effects of adding the PARP1/2 inhibitor olaparib (AZD2281, KU0059436) and the PTEN- activating phosphorylation inhibitor AZD454 using the same experimental protocols. All inhibitors, drugs or small molecules were obtained from SelleckChem (Houston TX) or from Sigma-Aldrich (St Louis MO) with the exception of ST-401, a gift from Dr. Nephi Stella (University of Washington Department of Pharmacology, Seattle WA).

**Whole exome sequencing library preparation and sequencing**

Whole exome sequencing was performed by targeted exon capture and sequencing on an Illumina HiSeq2500 or HiSeq 4000. Libraries for GBM4, GBM8 and 0827 sequencing were prepared using 150 - 200ng of DNA and the Illumina TruSeq Exome Preparation kit (Illumina, San Diego, CA) with the following modifications: amplified libraries were captured using the Roche Nimblgen SeqCap EZ Exome (v2) kit (Roche Sequencing and Life Sciences, Indianapolis,

IN) following the manufacturer's instructions, with the substitution of IDT xGEN dual index blocking oligos (Integrated DNA technologies, San Diego, CA ) for Nimblegen kit blockers. Hybridization of the resulting library to the probe set was at 47℃for 72 hrs. Capture washes were performed following the Nimblegen protocol on a Perkin Elmer Sciclone G3 NGSx workstation. Libraries were then finished by PCR amplification for 13 cycles using Kapa HiFi HotStart Polymerase.

GBM4, GBM8 and 0827 genomic libraries (1 sample/lane) were subjected to paired-end 75bp sequencing on an Illumina HiSeq 4000. For 0131, DNA was sonicated to ~200bp then subjected to KAPA A-tailing, end repair and adapter ligation followed by hybridization to a NimbleGen SeqCap EZ HGSC VCRome capture panel (Roche Sequencing and Life Sciences, Indianapolis, IN). The resulting 0131 library was then loaded on 2 lanes of an Illumina HiSeq2500 for sequencing. Both sets of raw sequence data were analyzed jointly. Primary data processing including quality control analysis and mapping using the nf-core sarek pipeline (Nextflow v 22.04.3; sarek v3.0)(Ewels et al., 2020). Read pre-processing, adapter trimming and quality filtering were carried out using fastp; both fastp and FastQC were used for quality control analysis (Chen et al., 2018). All samples passed QC using standard metrics and were aligned to reference sequence GATK.GRCCh37 with decoy sequences (human_g1k_v37_decoy.fasta) using bwa (Li and Durbin, 2010) and manufacturer-provided target capture regions specific for SeqCapEZ v2 and VCRome v2.1 capture libraries.

Freebayes, Mutect2, and strelka were used for somatic mutation calling against a publicly available panel of normals (Broad Institute: https://www.broadinstitute.org/genomics). Final mutation calls were based on a consensus of at least two of three callers, then annotated using Ensembl Variant Effect Predictor (https://grch37.ensembl.org/info/docs/tools/vep/index.html). Final results were summarized across tools and methods using Multiqc.

GSC data were compared with a TCGA Glioblastoma dataset (Brennan et al., 2013) and data from associated Cancer Genome Atlas analysis projects to include both *IDH*+ and *IDH* neomorphic-mutant samples: a total of 2,539 GBM tumors from seven different studies were included (see: cBioportal (https://www.cbioportal.org/) for 'CNS/Brain - Cancer Subtype Designation Glioblastoma multiforme' samples (accession date 7 April 2025). We also had access to, and drew selectively on, previously reported analyses of our GSC cultures (e.g., (Eyler et al., 2020; Toledo et al., 2015), and an ASCO-Tempus GBM dataset of >1,000 additional GBM patients/analyses

(https://www.asco.org/sites/new-www.asco.org/files/content-files/research-and-progress/documents/2020-Gliobloastoma-Data-Description.pdf)

Mutational Signatures Analysis was used to identify SNV/indel variants: mutect2 in the GATK toolkit (https://gatk.broadinstitute.org/hc/en-us/articles/360037593851-Mutect2) was used to analyze post-processed, combined .maf files. SigProfileAssignment (Díaz-Gay et al., 2023) was used to assign COSMIC GSC-specific mutation signatures (Alexandrov et al., 2020) as described at (https://cancer.sanger.ac.uk/signatures, Human Cancer Signatures v3.4 access date 25 April 2025). The following known artifactual signatures were excluded: single base substitution (SBS) signatures SBS27, SBS43, SBS45, SBS46, SBS47, SBS48, SBS49, SBS50, SBS51, SBS52, SBS53, SBS54, SBS55, SBS56, SBS57, SBS58, SBS59, SBS60, SBS95, together with the double base signature DBS14.

**Mitochondrial DNA variant detection**

Duplex DNA sequencing was performed as previously described (Kennedy et al., 2014) with mtDNA-specific modifications (Hoekstra et al., 2016; Sanchez-Contreras et al., 2021). In brief, Duplex-Seq adapters containing double-stranded unique sequence UMIs (IDT, Coralville IA) were ligated to sonicated DNA samples that had been end-repaired using the NEBNext Ultra End II Repair/dA-Tailing and Ligation kits (New England BioLabs, Ipswich, MA) following the manufacturer's instructions. We used qPCR to normalize mtDNA input amounts prior to sample amplification to incorporateTruSeq adapters (MWS13s: 5'-ACACTCTTTCCCTACACGACGC and MWS20: 5'-GTGACTGGAGTTCAGACGTGTGC, Illumina). Amplified libraries were then used for targeted capture by probes specific for human mtDNA (Integrated DNA Technologies, Coralville, IA), following the IDT xGen Lockdown protocol according to manufacturer instructions. The resulting libraries were indexed and sequenced using ~150–cycle paired-end reads (300-cycles total) on an Illumina HiSeq4000 to generate ~$20x10^6$ reads per sample.

Raw sequencing data were processed using an in-house bioinformatics pipeline (see: https://github.com/Kennedy-Lab-UW/Duplex-Seq-Pipeline; version 2.1.2 ) with default consensus-making parameters. In brief, post-consensus fastq files were aligned against the hg38 reference genome using bwa v0.7.17 (Li and Durbin, 2009) after masking seven small polynucleotide repeats to reduce alignment artifacts. Non-SNP variants (i.e., variants with allele fractions <40%) were BLAST-aligned against a database to identify and remove potential common contaminants and known pseudogenes from dog (canFam3), cow (bosTau9), nematode (ce11), mouse (mm10); rat (Rnor 6.0) and humans (hg38).

Variant reads that passed BLAST filtering were used with non-mutated/control reads to calculate the frequency of *de novo* events. A clonality cutoff of 1% (or a read depth of <100) was used to exclude variants with high heteroplasmy levels, prior to dividing allele read counts by the total number of reads at each mtDNA bp position. Variant calling was performed by VarDict-Java, with insertion-deletions (indels) left-aligned during variant calling. Unique variants were reported and plotted once per genome position and sample, with the frequency of *de novo* events quantified as described (Sanchez-Contreras et al., 2023). Variant protein coding changes were predicted using the Ensembl Variant Effect Predictor (VEP release 109, Feb 2023; see: https://grch37.ensembl.org/info/docs/tools/vep/index.html and McLaren et al., 2016). A more detailed description of this Duplex Sequencing pipeline can be found in (Sanchez-Contreras et al., 2021).

**Bulk RNA library generation and sequencing**

Bulk RNA gene expression analyses were performed using triplicate biological samples of 5x10e5 cells/GSC culture. RNA isolation, library prep and quality controls were performed following Seattle Genomics in-house protocols (https://www.seattlegenomics.com/). In brief, total RNA was extracted and purified using the miRNeasy micro kit (Qiagen,Germantown MD), the analyzed for quality on a Bioanalyzer prior to total RNA-seq library preparation using the rRNA depletion KAPA RNA HyperPrep Kit with RiboErase (HMR)(Roche Sequencing and Life Science, Indianapolis, IN). The resulting cDNA libraries were cleaned up and quantified on a Qubit using a dsDNA High Sensitivity Kit, then checked on an Agilent D100 ScreenTape Assay on a 4200 TapeStation System (Agilent, cat# 5067-5583) prior to pooling and NextSeq mid- and high output, 150 cycle sequencing (Illumina, San Diego, CA).

Raw (FASTQ) RNAseq files were analyzed using the nextflow (ver 22.04.3) pipeline (Di Tommaso et al., 2017), nf-core/rnaseq (version 3.7)(Ewels et al., 2020) and a bioinformatics workflow that integrated software tools designed RNA sequence analysis, further optimized to support parallel processing and minimize computational time. Read quality control analysis were performed using FastQC https://www.bioinformatics.babraham.ac.uk/projects/fastqc, with filtering of ribosomal RNA reads by SortMeRNA (Kopylova et al., 2012). Genomic contaminants were removed by BBSplit (BBMap - Bushnell B. - sourceforge.net/projects/bbmap/) prior to read quality assessment and adapter trimming using Trim Galore. STAR (Dobin et al., 2013) was used to align filtered reads to the human GRCh37 reference genome. SAMtools (Li et al., 2009) were used to sort and index alignments with Picard (‘Picard Toolkit 2019’ Broad Institute: https://broadinstitute.github.io/picard Broad Institute) to mark duplicate reads.

Expression was quantified using RSEM (Li and Dewey, 2011) and Salmon pseudo-alignment (Patro et al., 2017). Read processing, alignments and quality metrics were summarized using MultiQC (Ewels et al., 2016) prior to creating read coverage files using BEDTools (Quinlan and Hall, 2010) and bedGraphToBigWig from the UCSC Genome Browser toolkit (Kent et al., 2010; Quinlan and Hall, 2010). Read count normalization, QC analyses and differentially expressed genes were identified using the DESeq2 R package (Love et al., 2014), with a Benjamini-Hochberg adjusted p-value of 0.1 and log2 fold change of |1.3| set as thresholds to identify significantly differentially expressed genes.

**Single cell RNA sequencing and data analysis**

Single cell RNA sequencing was performed on aliquots of 20,000 cells/GSC culture taken on Day 0 of the time course sampling experiment (**Manuscript Figure 5A**) using the 10X Chromium Next GEM Single Cell v3.1 3' Reagent Kit and protocol (10X Genomics, Pleasanton, CA) according to manufacturer instructions. The resulting cDNA was captured and sequenced as described above without further enrichment using a standard 10X sequencing protocol (10X Genomics, Pleasanton, CA) on an Illumina NextSeq sequencer. In contrast to most single cell profiling gene expression analyses, mtDNA sequences were retained to allow expressed mtDNA variants to be identified and verified in Duplex mtDNA DNA sequencing data.

Gene-specific expression was identified in raw base call (BCL) files from Illumina sequencing that were demultiplexed into fastq files, then count matrix-mapped to a human hg19 reference genome using Cell Ranger (Zheng et al., 2017) with an 'expected cell number' = 20,000. Estimated cell counts ranged from 5906 - 7502/GSC culture, with a median detected gene number/cell of 2,059 - 2,695 and median UMI counts/cell of 5,802 - 8,309. Final expression counts were based on genes with ≥ 20 reads/cell in cells that expressed ≥ 200 genes, further qualified by UMI counts ≤ 25,000 and mitochondrial read percentages of ≤20%. These thresholds collectively identified 5618 - 7371 unique cells/GSC culture that individually expressed from 15477 - 16887 genes. Downstream analyses including data visualization, cleaning, geneset scoring etc. were performed using the Python scanpy package (version 1.10.4) (Wolf et al., 2018).

**Gene expression functional enrichment analyses**

Bulk RNA-seq analyses were performed using ssgsea (single sample gene set enrichment analysis, (Barbie et al., 2009) as implemented in the GSVA package in Bioconductor R version 4.2.1; GSVA version 1.46.0 (Hänzelmann et al., 2013). For single cell transcriptional profiling data, we applied the 'score_genes' function in scanpy (Python version 3.11.7, scanpy version

1.9.6)(Wolf et al., 2018) that reimplements Seurat ‘module scoring’ (Tirosh et al., 2016a). BulkRNA-seq enrichment scores (ES) represent normalized enrichment scores (NES) from ssgsea (Barbie et al., 2009) and scRNA-seq ES scores are the module scores as implemented in scanpy (version 1.9.6) as noted above, after Tirosh et al. (Tirosh et al., 2016b).

Expression enrichment in key glioma biology-related gene sets was observed for cell cycle phase distribution (Neftel et al., 2019); stemness; cancer hallmarks (Liberzon et al., 2015; Zhang et al., 2020); and glial cell surface markers. In addition, we looked for functional enrichment and previously defined gene expression signatures related to GBM subtypes and transcriptional cellular states (Wang et al., 2017; Al-Dalahmah et al., 2023; Garofano et al., 2021; Neftel et al., 2019; Venkataramani et al., 2022) using reference gene sets listed with their sources in **Table S8**.

GSC transcriptional states were compared with primary TCGA samples using a Functional Module States framework (Qin et al., 2022) that combined TCGA GBM and GSC culture data. RNASeq-based gene expression data were downloaded from TCGA GBM samples, with filtering to identify genes expressed in at least 5 samples of the profiled sample set (see:. https://gdc.cancer.gov/about-data/publications/pancanatlas). The distribution of log-transformed gene counts +1 from GSC cultures is similar to the distribution of log-transformed gene expression data +1 from the TCGA GBM sample set. Functional modules were used to assess activity: amino acid metabolism, apoptosis, carbohydrate metabolism, cell cycle, cell motility, cellular community - eukaryotes, cellular senescence, classical, energy metabolism, folding sorting and degradation, glycan biosynthesis and metabolism, lipid metabolism, membrane transport, metabolism of cofactors and vitamins, metabolism of other amino acids, nucleotide metabolism, DNA replication and repair, signal transduction, signaling molecules and interaction, transcription, translation, transport and catabolism, xenobiotics biodegradation and metabolism and the TP53 signaling pathway. We also included three gene sets to represent the GBM subtypes of classical, mesenchymal and proneural clusters.

Single sample gene set enrichment scoring (ssGSEA) was used for FMS module analyses, with ssGSEA scores for classical, mesenchymal, and proneural GBM gene sets used to identify three consensus clusters as a least ambiguous solution. These three resulting clusters reflect the TCGA sample subtypes of GBM ‘Classical’, ‘Mesenchymal’ and ‘Proneural’, to which GSC cultures were assigned to identify biological similarities between GSC cultures and TCGA samples.

GBM neurodevelopmental and metabolic subtypes were after Garofano et. al. (Garofano et al., 2021). To detect 'Neftel states' (Neftel et al., 2019), we computed six state scores: MES1, MES2, NPC1, NPC2, OPC and AC. NPC1 and NPC2 were merged into NPC, and MES1 and MES2 into MES by taking the maximum of each pair of values to represent NPC or MES, respectively. One state (MES, NPC, AC or OPC) was then assigned to each cell as defined by Neftel (Neftel et al., 2019). 'Undetermined' was assigned if the score for the second meta-module was higher than that of 10% of the cells mapped to this meta-module (as their top-scoring meta-module).

Recently identified gene expression signatures that enable cell-cell communication, electrical coupling and synaptic connectivity between tumor cells and with non-tumor cell types (Venkataramani et al., 2022; Venkatesh et al., 2019) as well as functional states that enable invasion and proliferation (Kim et al., 2024; Liu et al., 2024; Taylor et al., 2023) were extracted as reported, then used to search GSC bulk and scRNAseq data to capture important metabolic and functional state information and provide insight into GSC biology and predicted behavior.

**Proteomic and phosphoproteomic sample prep and profiling**

Mass spectrometry-based proteomic and phosphoproteomic profiling were used to characterize GSC cultures and their response to IR treatment. Pilot experiments were used to define IR dose-survival curves for all four GSC cultures, and to identify radiation doses and early sampling times that led to robust, early γ-H2AX (Ser139) phosphorylation with high cell survival. Proteomic analyses used 12 replicate T-25 flasks that were seeded with 2.5x10e6 cells/flask, allowed to recover for 24 hrs in a 5% oxygen atmosphere, then sealed to maintain a 5% oxygen atmosphere followed by irradiating 6 experimental flasks/culture with 1.0 Gy of X-radiation in parallel with 6 mock-treated control flask replicates. Radiation was delivered using a RS-2000 X-irradiator (Rad Source Technologies, Buford, GA) as detailed above. Samples for MS profiling analyses were prepared from irradiated or control cells at 1 hr after irradiation by brief Accutase treatment to generate single cell suspensions that were pelleted, flash frozen in liquid nitrogen and stored at -80°C.

Frozen cell pellets were resuspended in lysis buffer (8 M urea, 150 mM NaCl, and 100 mM Tris pH 8.2) prior to three rounds of sonication, with lysate protein concentrations determined by BCA assay. Proteins were then reduced with 5 mM dithiothreitol (DTT) for 30 min at 55°C, and alkylated with 15 mM iodoacetamide in the dark for 15 min at room temperature. The alkylation reaction was quenched by adding an additional 5 mM DTT, followed by incubation for 15 min at room temperature. Aliquots of 300μg of protein/sample were desalted and digested with trypsin using magnetic beads and a Kingfisher Flex (Thermo Fisher Scientific) magnetic particle processing robot following the R2-P1 protocol (Leutert et al., 2019). A peptide sample (25ug) was kept for total proteome analysis, with the remainder used for automated phosphopeptide enrichment on $Fe^{3+}$-IMAC magnetic beads as part of the above R2-P2 protocol.

**Mass spectrometry data acquisition and analysis**

Dried peptide/phosphopeptide samples were dissolved in 4% formic acid/3% acetonitrile for nLC- MS/MS analysis. Peptides were loaded onto a 100 μm ID × 3 cm precolumn packed with Reprosil C18 3 μm beads (Dr. Maisch GmbH), then separated by reverse- phase chromatography on a 100 μm ID × 30 cm analytical column packed with Reprosil C18 1.9 μm beads (Dr. Maisch GmbH) housed in a column heater set at 50°C. Peptides for total proteome analysis were separated by a 120 min gradient, and phosphopeptides by a 90 min gradient, of 7 to 28% acetonitrile in 0.125% formic acid on an Orbitrap Eclipse Tribrid Mass Spectrometer equipped with an Easy1200 nanoLC system (Thermo Fisher Scientific) operating in data-dependent acquisition mode.

For total proteome and phosphoproteome analyses, full MS scans were acquired from 375 to 1500 m/z at 120,000 resolution with a fill target of 4e5 ions and maximum injection time of 50 ms. A cycle time of 3s was chosen to select the most abundant ions on full MS scans for fragmentation using a 1.6 m/z precursor isolation window and beam- type collisional- activation dissociation (HCD) with 30% normalized collision energy. For total proteome analysis, MS/MS spectra were collected at 30,000 resolution with a fill target of 5e4 ions and maximum injection time of 54 ms. Fragmented precursors were dynamically excluded from selection for 60 s. Phosphopeptide MS/MS spectra were collected at 50,000 resolution with fill target of 1e5 ions and maximum injection time of 86 ms. Fragmented precursors were dynamically excluded from selection for 45 s.

The resulting MS/MS spectra were searched with MaxQuant(v.1.6.14) against the Uniprot human canonical + isoform protein sequence database. We used static modification of cysteine carbamidomethylation (57.021463 Da) and variable modification of methionine oxidation (15.994914 Da) for all searches, and variable modification of serine, threonine, and tyrosine phosphorylation (79.966331 Da) for phosphopeptide sample analyses. Trypsin/P was specified for protease digestion product identification, allowing for up to two missed cleavages. A target-decoy database search strategy was used to guide filtering and estimate FDRs. All data were filtered to 1% FDR at both peptide and protein levels with the 'match between runs' option enabled for a time window of 0.7 min to identify between-replicate matches. Proteins detected by at least two peptides - with one unique to the protein - were considered identified. For quantitative protein and phosphorylation site analysis, we used generated ‘proteinGroups.txt’ and ‘Phospho(STY)Sites.txt’ tables respectively after filtering off contaminants and reverse hits.

Bioinformatic analyses were performed using R (https://www.r-project.org/): phosphopeptide intensities were median-normalized, and phosphopeptide intensities were imputed with a shifted normal distribution of small values if they were completely missing in all 5 replicates in one biological condition, but present in other conditions and measured in 3 out of the 5 replicates. Statistically significantly changing phosphosites and proteins were determined using the limma package (doi: 10.18129/B9.bioc.limma), with filtering for Benjamini–Hochberg multiple-hypotheses-corrected p-values of <0.05 and absolute fold changes of |>1.5|. Protein and phosphoprotein abundances and statistics are summarized in **Table S11**.

GO enrichment analyses were performed using the whole human proteome as background. Fisher exact testing with Benjamini–Hochberg multiple- hypothesis correction at the protein level was used to filter for a FDR <0.02 to identify enriched terms. The annotation of proteins used Gene Ontology (GO) Biological Processes; GO Cellular Components; and the Kyoto Encyclopedia of Genes and Genomes (KEGG). Kinase-Substrate Enrichment Analysis (KSEA) was performed using the KSEA app (Wiredja et al., 2017), and protein motif enrichment analysis was performed using iceLogo (Colaert et al., 2009).

**Molecular barcoding to define population heterogeneity and track ionizing radiation response**

GSC-specific mtDNA variants and lentiviral randomized 8 bp DNA barcodes embedded in the untranslated 3’ region of an expressed lentiviral EGFP transgene (the pSMAL-CellTag-V1 system; Biddy et al., 2018) were used as molecular barcodes to identify and track individual GSC cells. The mtDNA variants were newly identified and characterized here by mtDNA-targeted Duplex sequencing analysis (Kennedy et al., 2013). The lentiviral CellTag-V1 plasmid library (pSMAL-CellTag-V1), used to transduce and label GSC cultures contained ~20,000 unique molecular barcodes (Addgene Library 115643), was kindly provided by Dr. Anoop Patel, UW-Seattle Neurosurgery (now at Duke University, Durham NC). The lentiviral plasmid library was amplified in *E. coli,* purified by CsCl ultracentrifugation, and packaged following a Morris Lab protocol (Biddy et al., 2018) detailed at: (https://www.protocols.io/view/single-cell-mapping-of-lineage-and-identity-via-ce-kxygxm33wl8j/v8).

Pilot experiments were performed to identify the best protocol to transduce an average of at least 3 barcodes/cell, as estimated from a multiplicity of infection (MOI) versus resulting EGFP-positive cell fractions that were quantified by flow cytometry. Prior to lentiviral library packaging, we PCR-amplified the pSMAL-CellTag-V1 plasmid barcode region for Illumina short read sequencing to define a ‘white list’ of starting library barcodes. In brief, a common primer pair flanking the CellTag V1 barcode region was used to generate a 134 bp amplicon from starting lentiviral vector plasmids. The same primers also allowed the amplification of the barcode region of integrated copies of the CellTag V1 vector from GSC cultures. Primer sequences were:

**Read2_CellTag_R1_primer:**

5’ - GTCTCGTGGGCTCGGAGATGTGTATAAGAGACAGGATCTCAAATCCCTCGGAAG - 3’, and

**Read1_GFP_primer:**

5’ - TCCCTAGACGACGCTCTTCCGATCTGGCATGGACGAGCTGTACAAGTAA - 3’.

Resulting lentiviral CellTag-V1 amplicon DNA was UMI-indexed prior to Illumina short read NGS sequencing using variants of workflows developed by the Morris (Biddy et al. 2018; https://github.com/morris-lab) and Trapnell Labs with help from Jose McFaline, UW Genome Sciences, Seattle WA and Columbia University). CellTag barcodes were identified in fastq files extracted from raw NGS paired end reads using bcl2fastq, using the primer pairs as seed sequences. In brief, Read2 (3’ to 5’) covered the 8 base CellTag sequence, with Read1 (5’ to 3’) providing an overlapping segment of the CellTag 3’ end sequence.

Total reads from GSC cultures transduced with an average of 3 vectors/cell ranged from 394,365 - 1,975,777, with a median of 718,887 reads/sample. Base quality was checked using FASTQC (https://www.bioinformatics.babraham.ac.uk/projects/fastqc/), followed by use of a customized script to identify CellTags from the paired reads as follows: First, nucleotides with Q-scores <30 at the 3’ end of Read1 (R1) were trimmed, and the complement of Read2 (R2) was generated to provide a 5’ to 3’ sequence to use as a search string to identify the CellTag-V1 barcode region sequence motif GGT(********)GAATT. A parallel search was performed in R1 sequence data using GGT(*****) as a seed to search for the last 11 nucleotides of R1 that overlapped the CellTag barcode region.

CellTags identified in R1 and R2 were then compared to identify the subset that unambiguously identified identical, unique tag sequences: these are summarized in **Table S9**, where the average proportion of reads specifying a unique tag sequence was 88.7% for GBM4, 88.6% for GBM8, 87.2% for 0131 and 87.6% for 0827. The frequency of individual, unique sequence tag reads was calculated from the number of reads for each unique tag sequence divided by the total number of reads that identified CellTags in each sample. In order to perform meaningful downstream analysis, we selected the subset of these unique tags in each GSC culture with ≥10 reads in at least one sample of a given culture. This thresholding strategy captured an average of 96% of all detected CellTags, and 20% of all unique CellTags.

The mitotic stability of expressed CellTag lentiviral barcodes was determined by transducing 500,000 cells/GSC culture, then flow-sorting the transduced population to generate GSC cultures of 200,000 EGFP+ cells/GSC culture that were ≥95% EGFP+. These EGFP+ cell populations were then serially passaged with sampling at Days 3, 21 and 49 to determine the fraction of cells that retained EGFP expression. All four GSC cultures remained >95% EGFP+ over 49 days, which indicated that the expressed CellTag system and EGFP expression were sufficiently stable to permit post-IR time course sampling over up to 50 days.

**Cellular trajectories inferred from CellTag barcode and scRNAseq data**

A time point sampling experiment was designed that used mtDNA variants and CellTag molecular barcodes to define GSC culture cellular heterogeneity, and then identify gene expression programs that might favor cell proliferation or loss over time and after IR treatment (**Figure 5A**). This protocol was initiated by flow-sorting 20,000 EGFP+cells/GSC culture that were expanded to ≥2x10e6 cells/culture that were used to seed all experimental arms on Day 0 and to provide initial characterization samples (**Figure 5A**). Experimental arms consisted of: a control (untreated) arm, an IR-treated arm, and a 'reduced complexity' arm consisting of triplicate cultures of 40,000 cells each to mimic the effect of an 80% reduction in viability in the IR-treated arm though without radiation exposure. IR doses of 1.6 - 2.5 Gy, determined in pilot experiments, were used in +IR treatment arms to reduce cell populations by an estimated ~80%; within-experiment survivals were quantified by CFE scored Day 21 after IR treatment.

Day 0 characterization samples were used for bulk (triplicate samples of 500,000 cells/GSC culture) and single cell (20,000 cells/culture) RNA sequencing; CellTag barcode amplicon sequencing; and mtDNA variant detection by Duplex DNA sequencing (the latter two assays used 200,000 cells/assay/GSC culture). Amplicon sequencing of lentiviral CellTag barcodes identified ~6000 - 8000 unique CellTag barcodes/GSC culture, with many barcodes identified in all four cultures. In 0131 these 'overlap' tags represented >50% of all unique tags, reflecting the substantially higher integrated lentiviral vector copy number/cell in this culture.

Cultures were propagated after Day 0 with sampling of 200,000 cells/culture on Day 7 (all 4 cultures), Day 12 (0131 and 0827), Day 15 (GBM8), and Days 20, 3 and 40 (all 4 cultures). Different sampling timepoints were used to allow sufficient growth between sampling timepoints to ensure that no sample exceeded 20% of the total cell population at a given time point in order to minimize sampling-induced ‘bottle-necking’ of population complexity. Reduced complexity triplicate cultures were sampled on Days 7, 12 or 15, 20, 30 and 40 days as above. We were able to sample from one to three replicate cultures for each GSC culture and time point where the missing cultures were lost due to fungal contamination. Lentiviral barcode and mtDNA variant sequencing data were generated from final sample sets of 17 to 26 timed samples/GSC culture. These identified 2716 (GBM4) to 5086 (0827) unique lentiviral barcodes and 84 (0131) to 142 (GBM4) mtDNA variants/GSC culture.

CellTag and mtDNA variant barcode trajectories were established by identifying barcodes present in Day 0 samples that could be tracked across additional time points. Lentiviral barcode read frequencies normalized to Day 0 samples were used to construct Day ($T_n$ /$T_0$) read count ratios that were log-transformed prior to K-means clustering using the Python ‘sklearn’ package. A similar general strategy was used to identify mtDNA variants present at Day 0 and in samples from additional time points and plot changes in their corrected frequencies over 40 days.

Five clusters of barcodes that displayed similar trajectories were identified in each GSC culture. Barcode clusters were further analyzed to identify trajectories with significant enrichment or depletion as a function of time and/or IR treatment. We first imputed a $T_n$ /$T_0$ ratio of $1x10^{-6}$ for any sample lacking a time point-specific measurement, then-subtracted the ratio of reads in the +IR experimental arm from the control (ctrl) arm sample at the same time point. Barcode read ratios at Day 40 were used to identify potentially enriched (ratios >2 of enriched/ctrl sample) and depleted (ratios <0.5, depleted versus ctrl sample) barcodes that were Wilcoxon tested to identify barcode subsets that displayed a significant change in frequency under either alternative hypothesis (>/enriched vs. </depleted).

Single cell sequencing on the 10X Chromium controller platform allowed us to capture expressed CellTag barcodes and mtDNA variants in conjunction with cell-specific UMI cell indexing. These data together allowed us to identify single cells and link their trajectories and their gene expression programs. This was done in two steps. In brief, we first identified CellTag barcodes in unmapped BAM file data using as a reference hg19. Barcode tag reads with > 20% ambiguous nucleotide calls were excluded first, prior to searching 10X scRNAseq data for the lentiviral CellTag-V1 sequence motif 'GGT[ACTG]{8}GAAT'. In order to track robustly expressed and adequately sampled expressed CellTag barcodes, we selected barcodes with >10 reads in at least one condition. CellTag barcodes that were enriched, depleted or unchanged over time were then identified by expression trajectory K-means clustering based on time course expressed variant $T_n$ /$T_0$ ratios. As individual GSC cells might contain different molecular barcodes, we identified cells as enriched or depleted only when >50% of the unique tags corresponded to that putative trajectory. Cells with both 50% enriched and 50% depleted tags were excluded as ambiguous.

Putative enriched or depleted cells were displayed as a UMAP of all single cells using Scanpy (Wolf et al., 2018). Differential gene expression between enriched and depleted cells was assessed using the function ‘scanpy.tl.rank_genes_groups’, and assessed by use of a Wilcoxon rank sum test. Commonly overexpressed genes or down-regulated expressed genes for different cultures were selected, and visualized using the ‘scanpy.pl.dotplot’ function.

Expressed CellTags that showed enrichment or depletion in IR-treated versus control samples were further assessed using the one-sided Wilcoxon test implemented in the ‘scipy.stats’ python library. A ratio of <0.5 or >2 between IR/control (Ctrl) samples on Day 40 with a threshold p-value of <0.1 was used to define the depleted or enriched tags associated with IR treatment. In order to identify genes and gene expression programs that might enable preferential cell enrichment or depletion over time and after IR treatment, we used the ‘scanpy.tl.rank_genes_group’ function together with a Wilcoxon rank-sum test to assess the significance of differences between groups. The ‘scanpy.pl.dotplot’ function was used to visualize the top shared differentially expressed genes identified between cell groups.

**Materials, Data and Code availability**

The GSC cultures reported here are available upon request and completion of a related Materials Transfer Agreement. Contact the communicating author for additional information.

**DNA/RNA sequencing data**

These data and records are archived under Bioproject PRJNA1036631 (https://www.ncbi.nlm.nih.gov/sra/PRJNA1036631) under the following submission records: whole exome sequencing data, SRA SUB15419289; Duplex mtDNA sequencing data, SRX24964902-SXR24964933; bulk RNAseq data, SRA SUB15367655 (raw reads) and GEO GSE303069 (processed data); single-cell RNA-seq data,GSE303662; and CellTag molecular barcode amplicon sequencing data, GEO  GSE302554.

**MS-MS proteomic data:** Mass spectrometry proteomics and phosphoproteomics data have been deposited with dataset identifier PXD035886 to the ProteomeXchange Consortium via the PRIDE Partner Repository.

**Code availability.**

In-house developed bioinformatics pipelines and code can be found at the following Github repositories:

For version 2.1.2 of whole exome and mtDNA Duplex data processing pipelines and code, see:

https://github.com/Kennedy-Lab-UW/Duplex-Seq-Pipeline

For gene expression and amplicon bar code analysis pipelines and code, see:

https://github.com/IlyaLab/GSC_culture_characterization

**Figure S1: GSC dose-response profiles for ionizing radiation, temozolomide and additional small molecule treatments.**

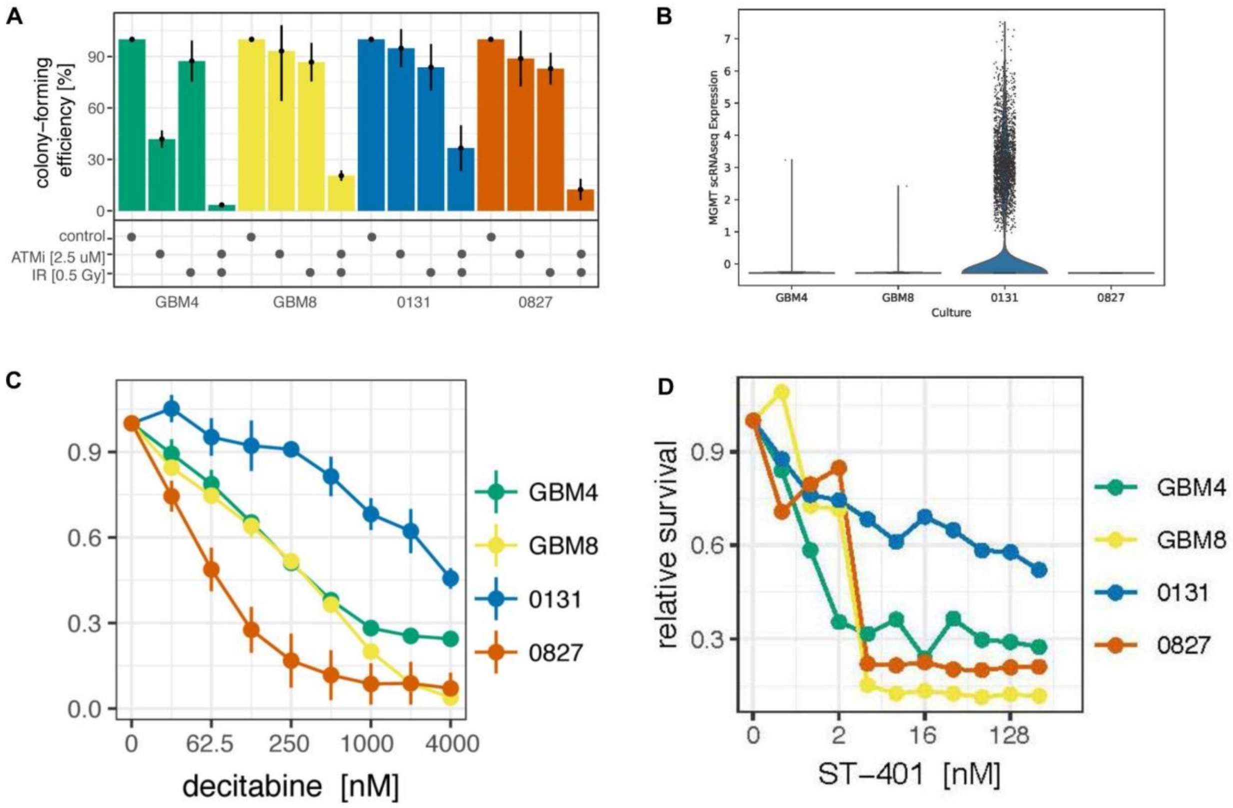


**Figure S1: GSC dose-response profiles for ionizing radiation, temozolomide and additional small molecules.** (A) CFE of GSC cultures when pre-treated with ATM kinase inhibitor (ATMi) KU-60019 prior to radiation with 0.5 Gy X-rays. (B) GSC MGMT expression quantified by scRNAseq analysis. These results are consistent with prior MGMT expression data from bulk RNAseq and Western blot analyses. (C) GSC fractional survival as a function of decitabine dose, a DNA demethylating agent used to reactive MGMT expression. (D) Potent GSC killing by nanomolar ST-401, a novel, brain-penetrant microtubule-targeting small molecule active in a mouse RCAS/tv-a *PDGFB* glioma model (see Horne et al., 2021; Vicente et al., 2024).

**Figure S2. GSC cultures are aneuploid with numerous numerical and structural chromosome abnormalities.**

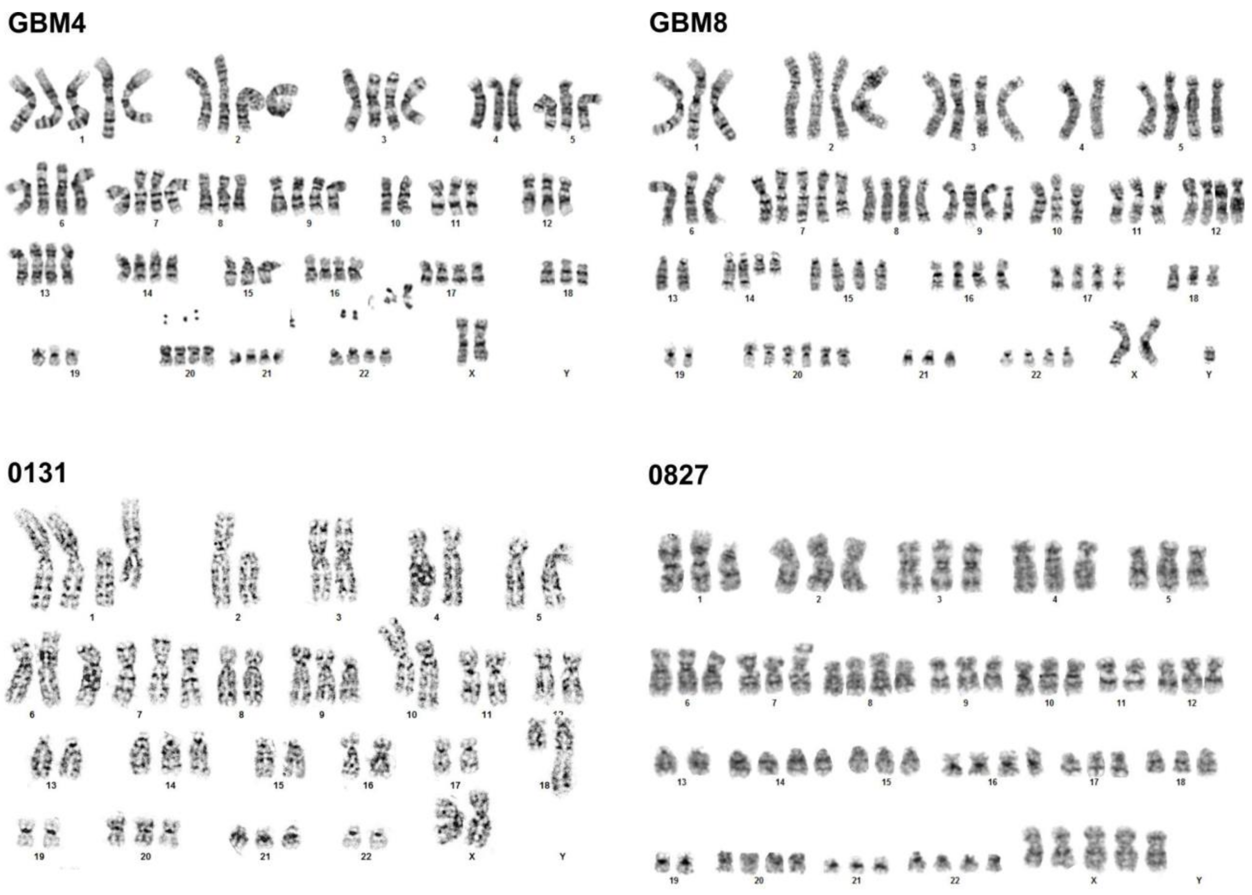


**Figure S2. GSC cultures are aneuploid with numerous numerical and structural chromosome abnormalities.** Representative G-banded metaphase karyotypes reveal aneuploid cultures with multiple, diverse numerical and structural abnormalities. Metaphase chromosome numbers ranged from 51 - 86/metaphase. GSC 0131 was near-diploid with several trisomic chromosomes; 0827 was near-triploid with chromosome-specific tri-, tetra- and pentasomies; and GBM4 and GBM8 were both near-tetraploid. Other recurrent high grade glioma chromosome abnormalities were observed in all cultures, together with marker chromosomes of unknown origins. The small unassigned chromosome in GBM8 was subsequently shown to be a non-Y-derived marker (additional results not shown; see Results and Methods for additional detail).

**Figure S3: GSC mtDNA mutational spectrum and subclonal structure identified by Duplex DNA sequencing.**

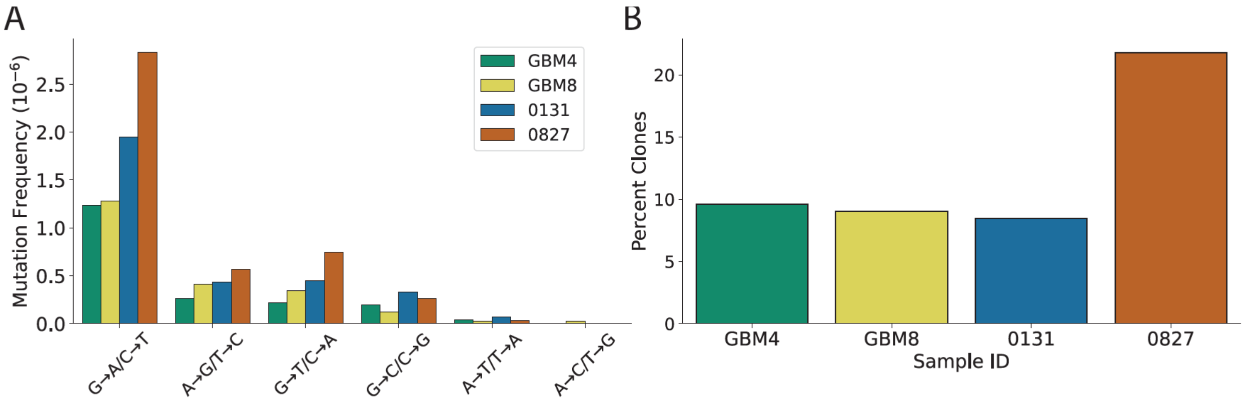


**Figure S3: GSC mtDNA mutational spectrum and subclonal structure identified by Duplex DNA sequencing. (A**) Mutational spectrum of GSC culture mtDNA displays a strong bias for G>A/C>T transitions, as previously reported, indicating that GSC culture conditions do not generate mutational artifacts. **(B)** Percentage of clonal variants (defined as >=2 supporting reads), where 0827 exhibits a much higher fraction of clones compared to the other three cultures.

**Figure S4: GSC mtDNA DNA base substitution variant (SNV) frequency, spectrum and subclonal composition as a function of time and IR treatment.**

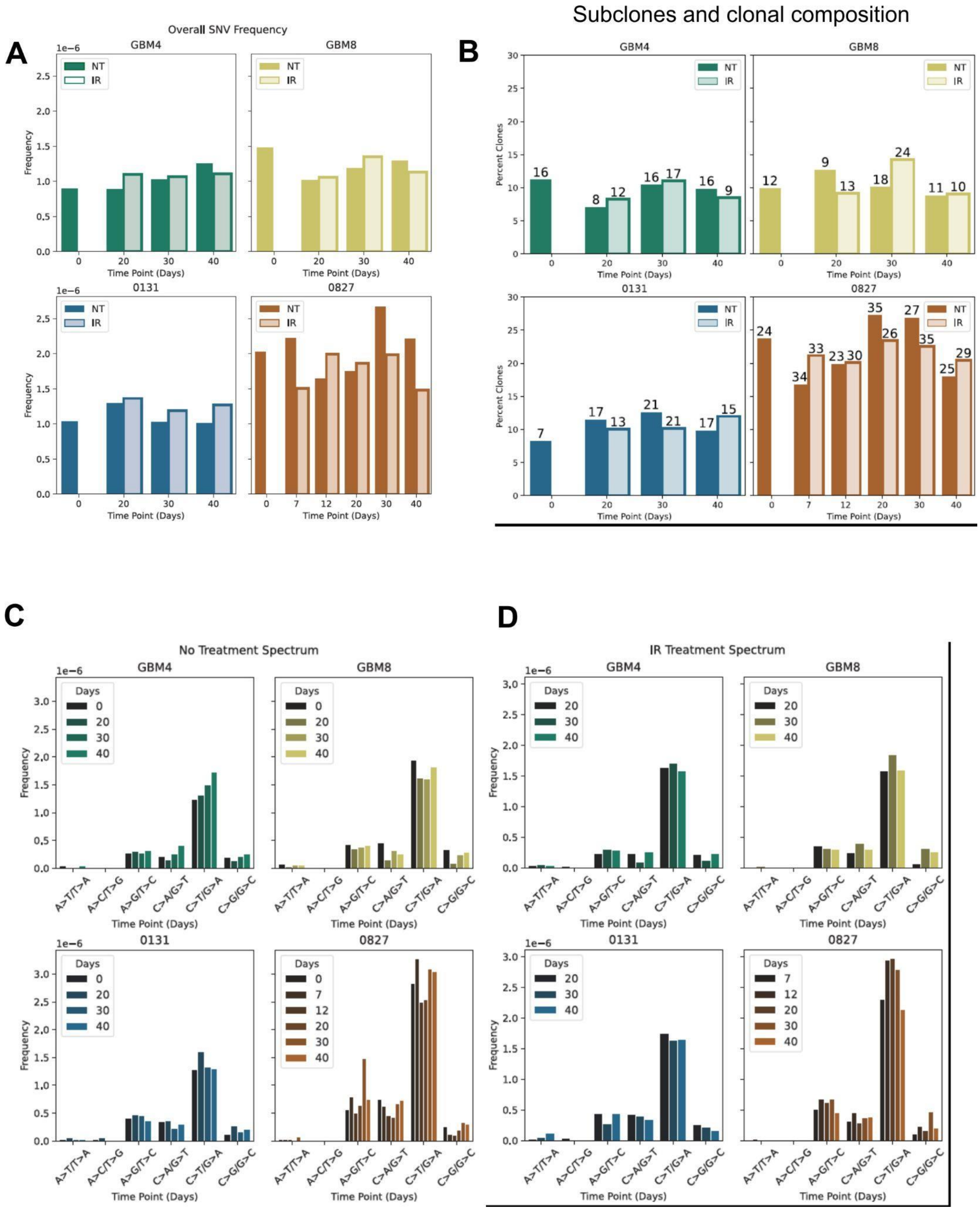

**Figure S4: GSC mtDNA DNA base substitution variant (SNV) frequency, spectrum and subclonal composition as a function of time and IR treatment. (A)** SNV frequency from time point samples taken over 40 days after IR treatment (NT) versus controls. **(B)** Percent of SNVs that are clonal (defined as >2 supporting duplex reads) in time point samples where the number above each bar indicates the number of clones detected. **(C, D)** Spectrum of mtDNA mutations and frequency of SNVs in IR-treated versus no-treatment controls revealed no significant change over the 40 day experimental time course

**Figure S5. GSC culture cell cycling metrics.**

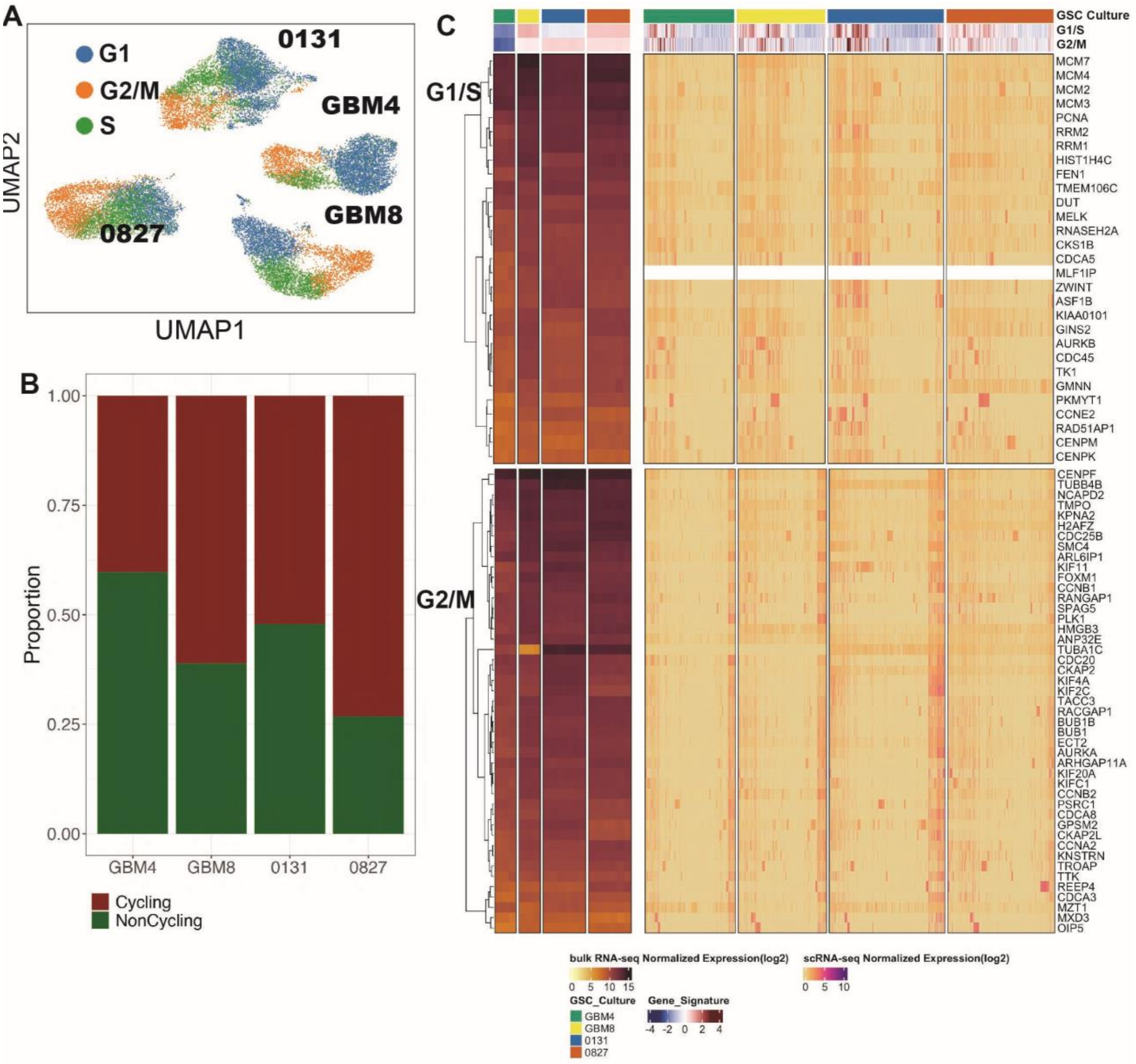


**Figure S5. GSC culture cell cycling metrics.** (**A**) Cell-cycle phases assigned based on G1/S and G2/M phase module enrichment scores from scRNA-seq in GSC cultures (**B**) Barplot depicting the proportion of cycling and noncycling cells (G2/M and S are cycling, G1 is noncycling). (**C**) Geneset enrichment scores for cell cycle-associated gene signatures (G1/S, G2/M) in (left) bulk RNA-seq and (right) scRNA-seq datasets (top-level heatmap) and per-gene level expression estimates for each bulk RNA-seq replicate sample (left) or single-cell (right) for each gene in the respective cell cycle signatures.

**Figure S6: GSC cultures exhibit transcriptionally-defined GBM cellular states.**

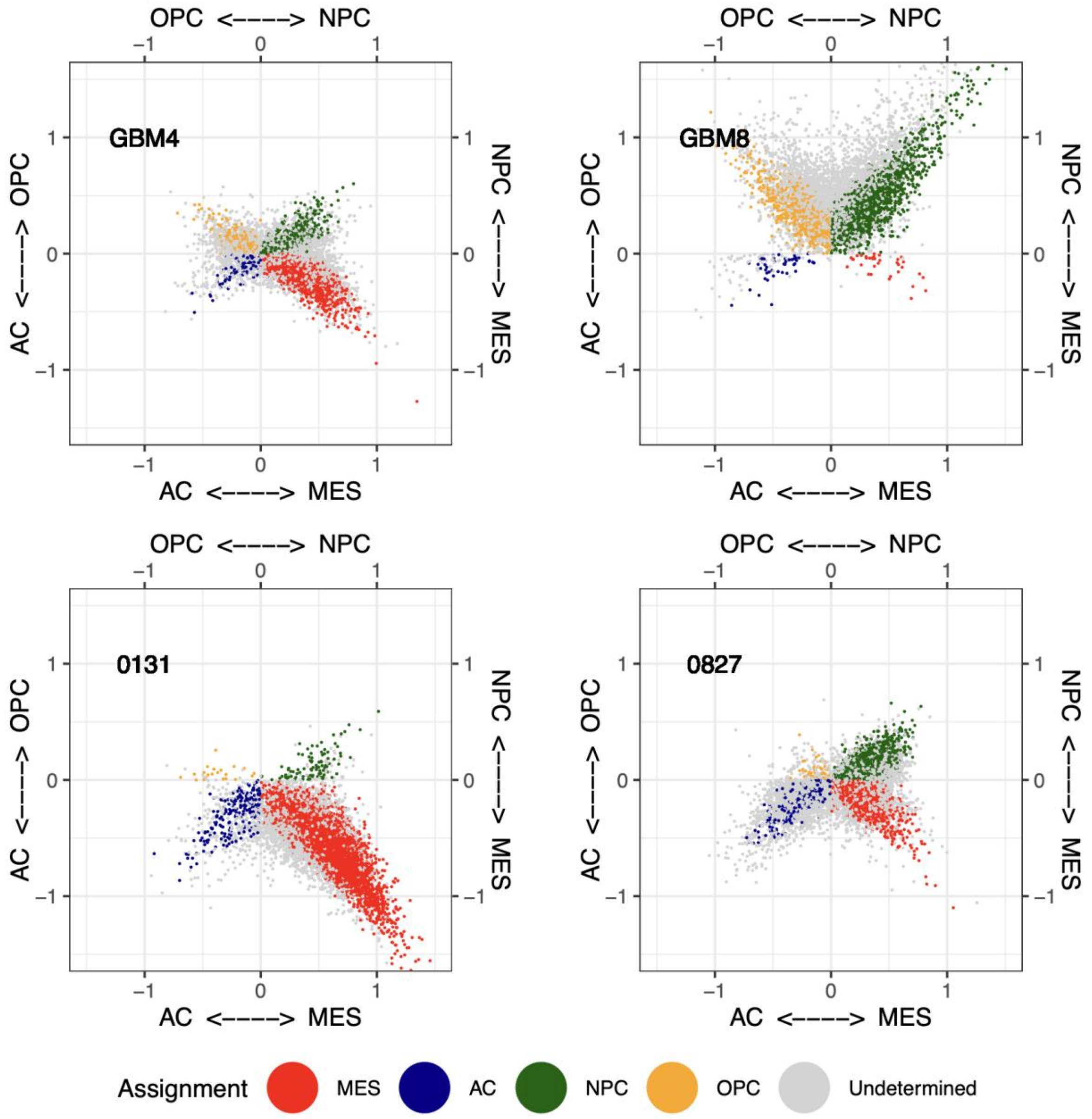

**Figure S6: GSC cultures exhibit transcriptionally-defined GBM cellular states.** Two-dimensional representation of single cell states defined by Neftel (Neftel et al., 2019), visualized using Scpubr tools (Blanco-Carmona, E. (2022) bioRxiv doi:10.1101/2022.02.28.482303). Each quadrant corresponds to one of the meta-modules (AC, MES, NPC or OPC), where the MES score is the maximum of MES1 and MES2 scores, and the NPC score is the maximum of NPC1 and NPC2 scores). The position of the dots (single cells) shows relative scores for the meta-modules). ‘Undetermined’, representing cells that could not be definitively assigned to any particular meta module, were common in all four GSC cultures and ranged from 31.9% (0131) to 59.4% (0827) of all cells analyzed (see Table S6 for distributions).

**Figure S7: GSC culture expression of markers of stemness.**

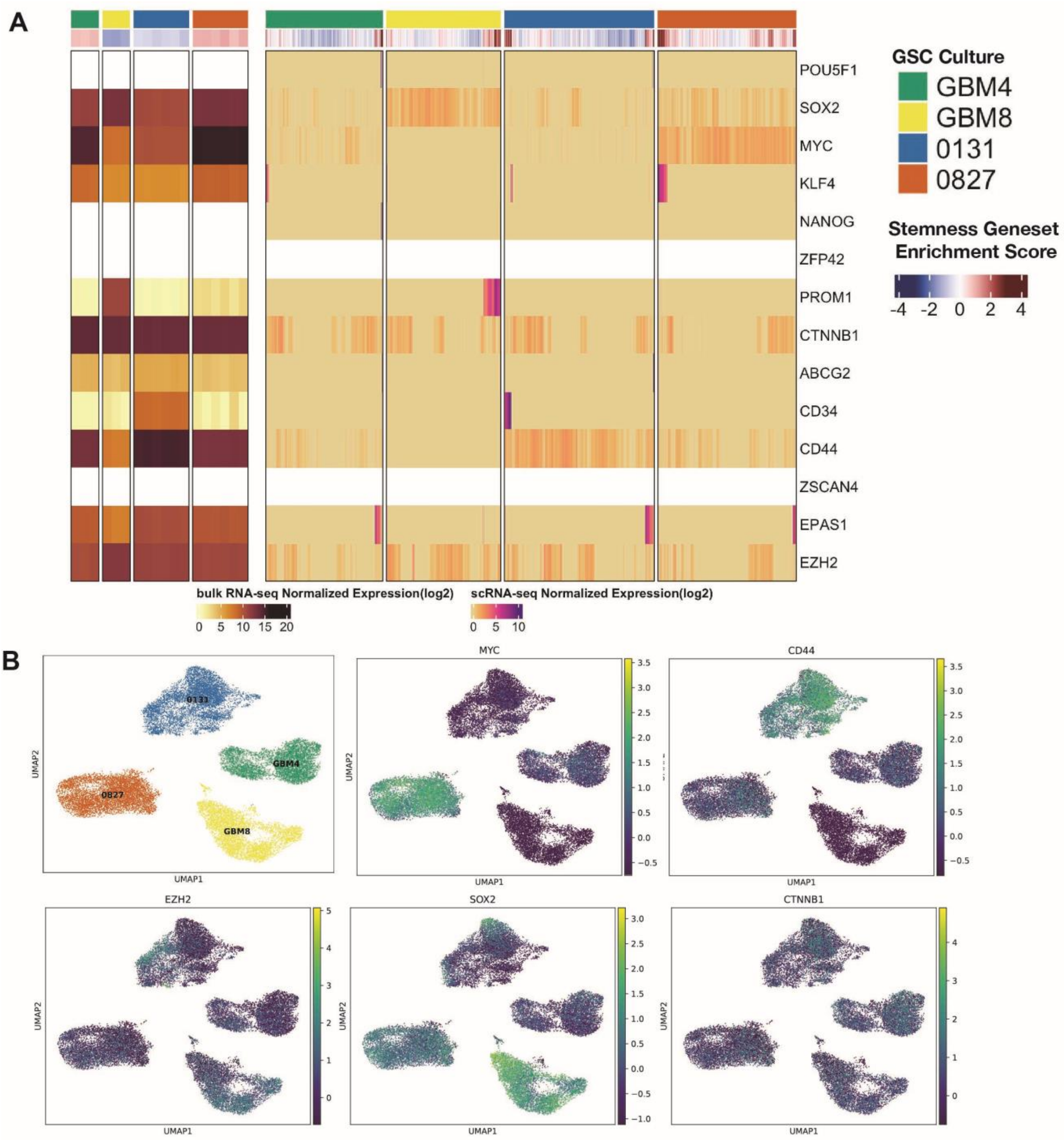


**Figure S7: GSC culture expression of markers of stemness.** (A) Expression of known stem cell markers as defined by (Malta et al., 2018) in (left) bulk and (right) single cell RNA-seq. The heatmap represents the quantified mRNA expression estimate for each stemness marker in the indicated signature. Genes not quantified on either platform are depicted in white. The annotation track (upper) represents geneset enrichment scores for the set of stemness-associated genes (STEMNESS). (B) UMAP representation of expressed stem cell signature genes in single cells across the four GSC cultures in the present study.

**Figure S8: Expression of GSC surface markers and key functional biomarkers.**

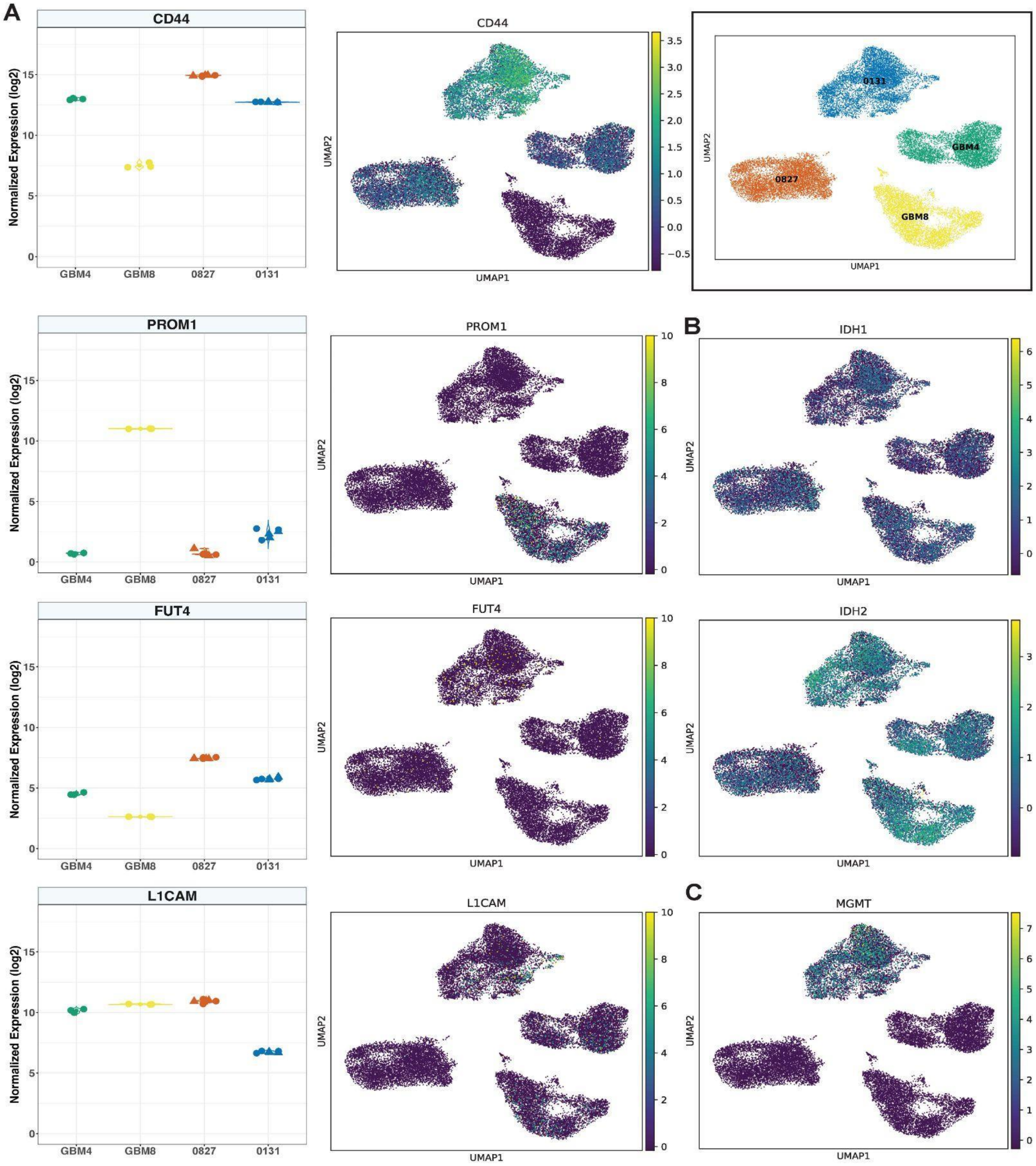

**Figure S8: Expression of GSC surface markers and key functional biomarkers. (A)** Expression of previously reported GSC-specific cell surface markers CD44, CD133/PROM1, L1CAM and FUT4 in bulk RNA-seq samples (left column) and at the single cell level (right column) in each culture. Expression of key GBM tumor biology genes and biomarkers **(B)** IDH1/2 and **(C)** MGMT at the single-cell level across all four cultures. GSC culture assignments are indicated in the top-right corner across all UMAP panels in the figure (inset). See Table S8 for additional information.

**Figure S9: GSC 10X scRNA sequencing data prior to and after filtering.**

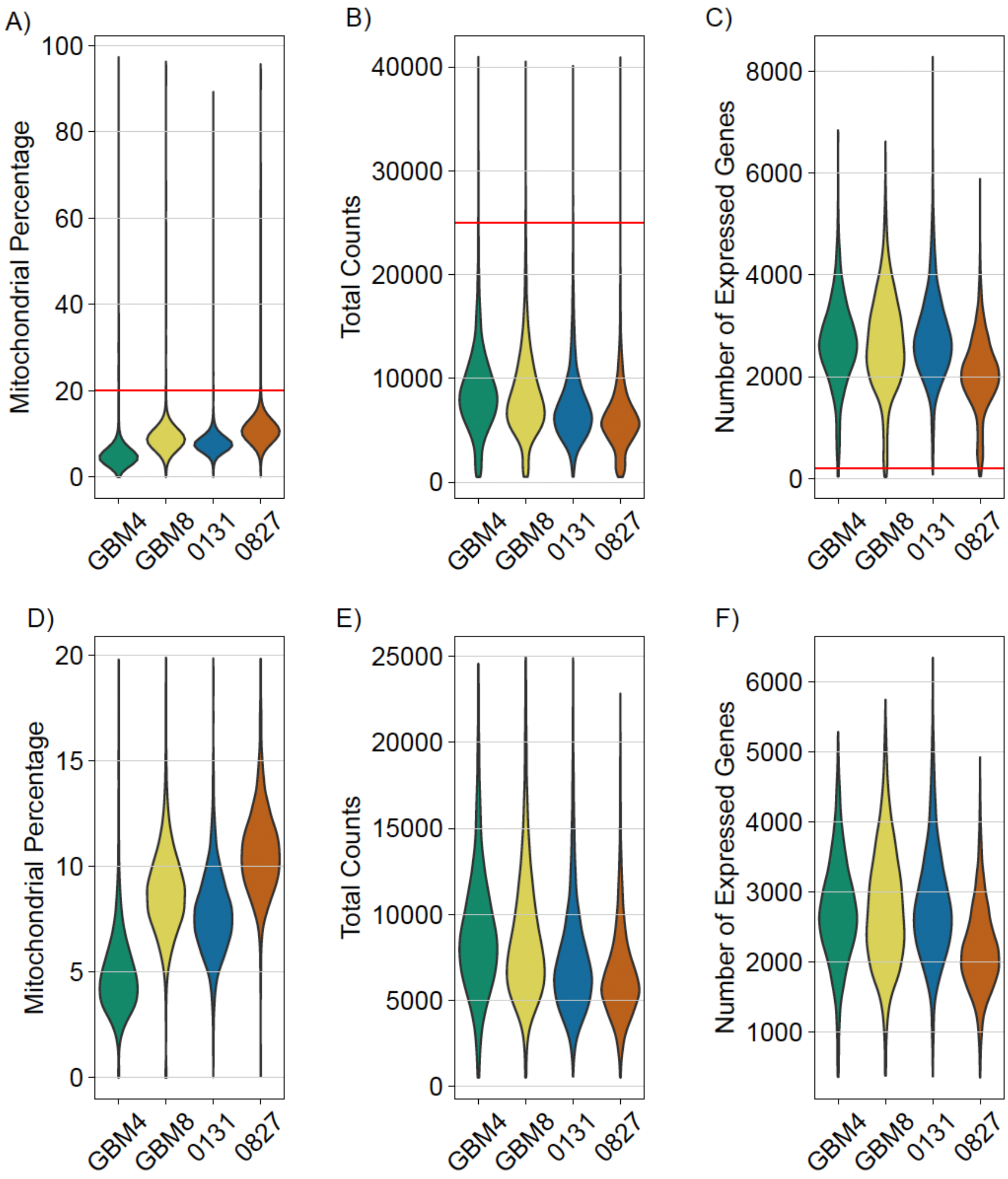


**Figure S9: GSC 10X scRNA sequencing data prior to and after filtering.** Mitochondrial percentages, total counts and number of expressed genes identified in 10X scRNA seq data prior to (top row, panels **A-C,** with thresholds used for filtering marked by the red horizontal line) and (bottom row, panels **D-F**) after filtering.

**Figure S10. Comparison of GSC and TCGA-GBM samples using Functional Module states scores based on gene expression.**

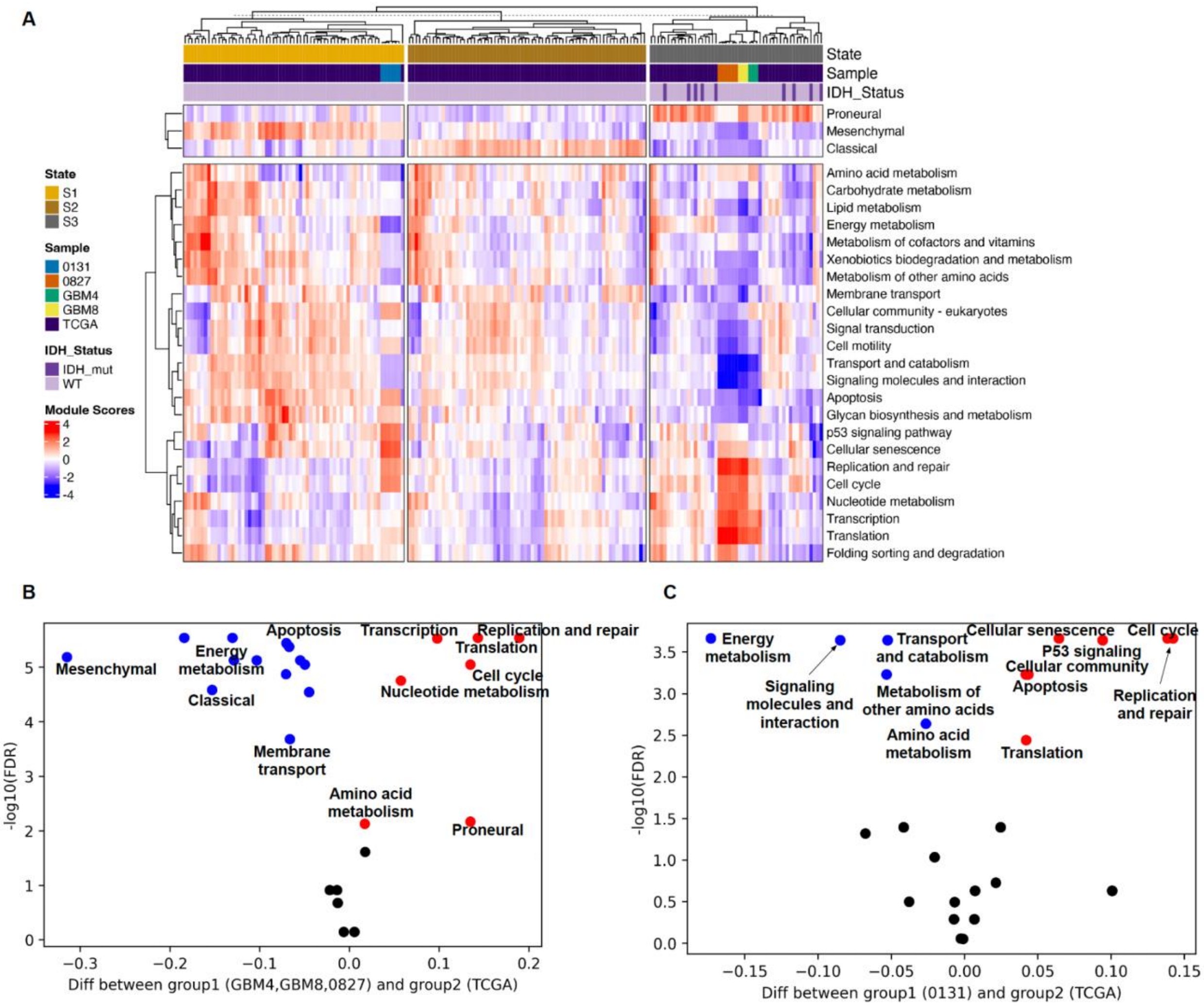


**Figure S10. Comparison of GSC and TCGA-GBM samples using functional module state scores based on gene expression. (A)** Three clusters identified by the module scores from GBM subtype related signature gene sets from the combined datasets of GSC cultures and TCGA-GBM samples. Functional modules scores were illustrated in the lower panel. GBM4, GBM8 and 0827 are clustered in a group with higher proneural score, and 0131 was clustered in another group with higher mesenchymal score (single sample gene enrichment score). **(B)** Comparison between the GSC cultures (GBM4, GBM8, 0827) and TCGA-GBM gene expression in the FMStates space. **(C)** Comparison between the GSC cultures (0131) and TCGA GBM gene expression in the FMStates space. A ranksum test was used followed by Benjamani-Hochberg adjustment. Red and blue colored dots represent modules which show a significant difference between the two compared groups (FDR < 0.01). Blue: Lower scores (activity) in GSC cultures; Red: higher scores (activities) in GSC cultures.

**Figure S11: Additional detail on GSC proteomic and phosphoproteomic characterizations including DNA metabolic protein complexes.**

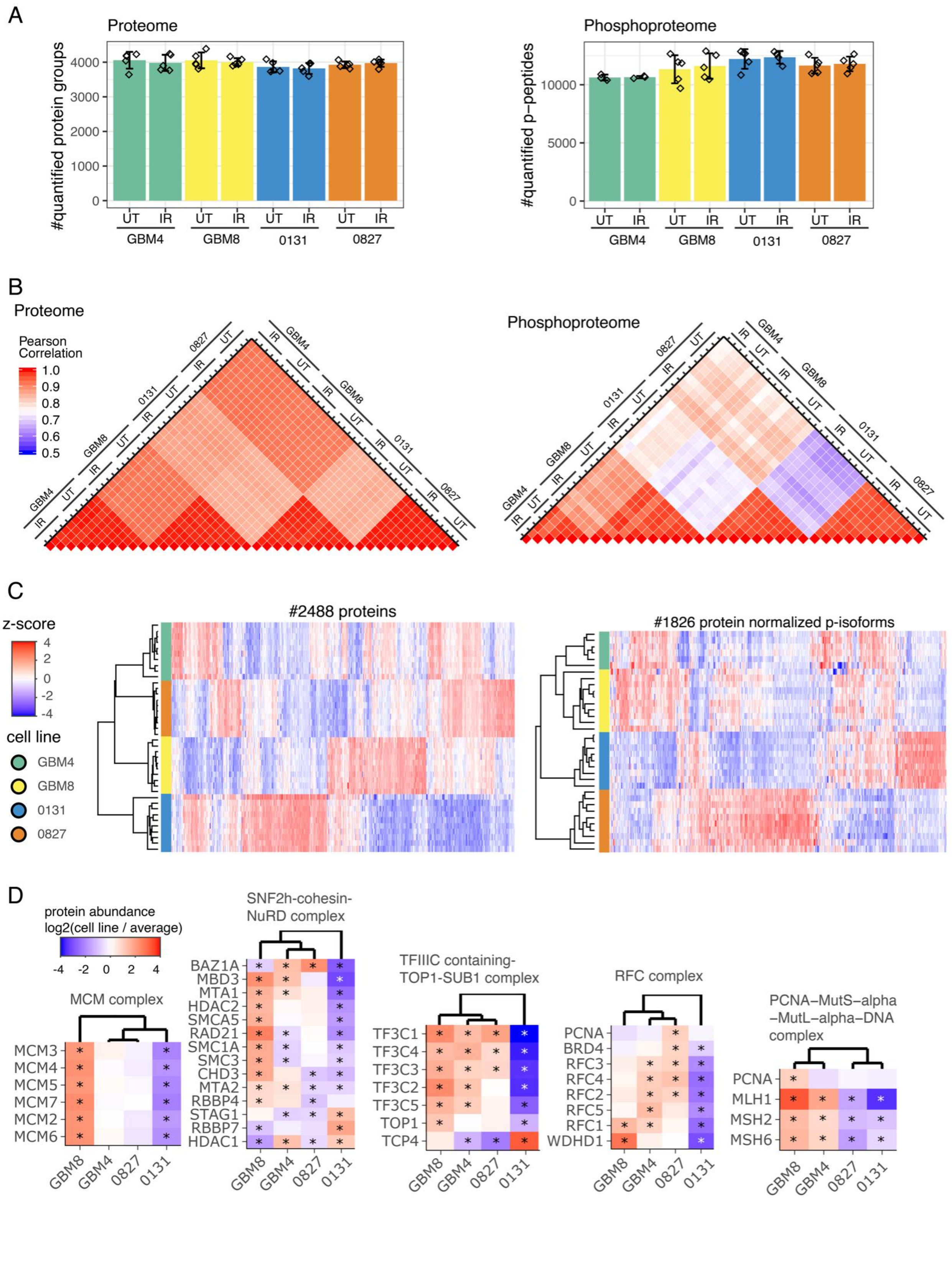

**Figure S11: Additional detail on GSC proteomic and phosphoproteomic characterizations including DNA metabolic protein complexes.** (A) Number of identified proteins and phosphorylated peptides in the different samples. (B) Heatmaps of Pearson correlation coefficients for protein and phosphopeptide quantifications. (C) Heatmaps of unsupervised hierarchical clustering of z-scored proteins and phosphosites that showed significant regulation in at least one condition. (D) Heatmaps of log2 fold changes (specific GSC culture versus average expression in all four cultures) of proteins associated with protein complexes involved in DNA metabolic processes that have been identified to be regulated in at least one cell line (* adjusted p-value < 0.05 and absolute fold change >2).

**Figure S12: GSC proteomic and phosphoproteomic response to ionizing radiation.**

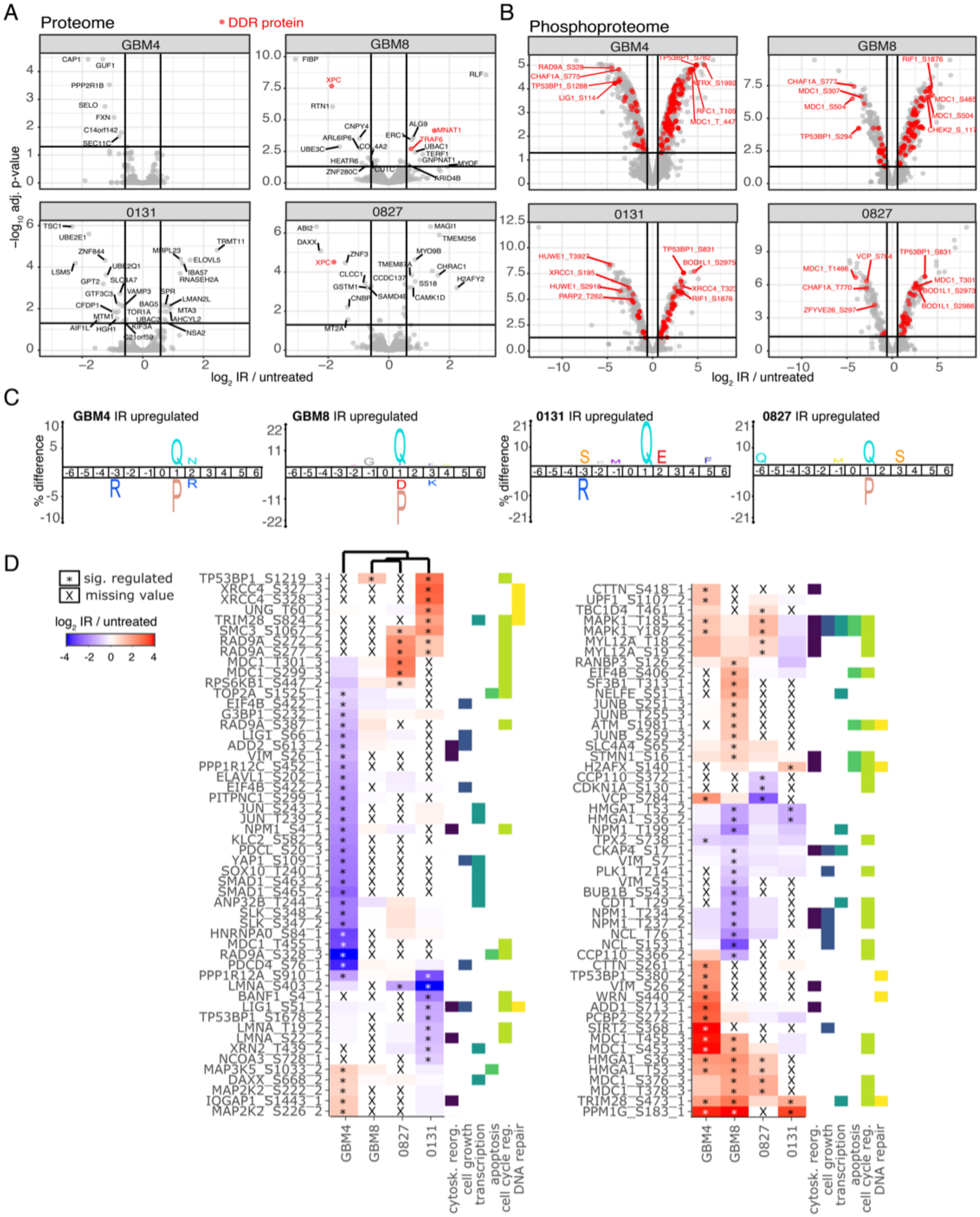

**Figure S12: GSC proteomic and phosphoproteomic response to ionizing radiation.** (A) Volcano plot depicting regulated proteins upon IR in the four GSC-cultures. Proteins associated with the DNA damage response are indicated in red. (B) Volcano plot depicting regulated phosphosites upon IR in the four GSC-cultures. Top 4 down and upregulated phosphosites in each cell line are indicated. Phosphosites on proteins associated with the DNA damage response are indicated in red. (C) iceLogo plot showing the enrichment and depletion of amino acids surrounding phosphorylation sites that are upregulated upon IR in the different GSC-cultures. (D) Heat map of phosphorylation sites with a known function as reported by Phosphosite Plus (Hornbeck et al., 2019).

**Figure S13: CellTag labeling and distribution in GSC cultures.**

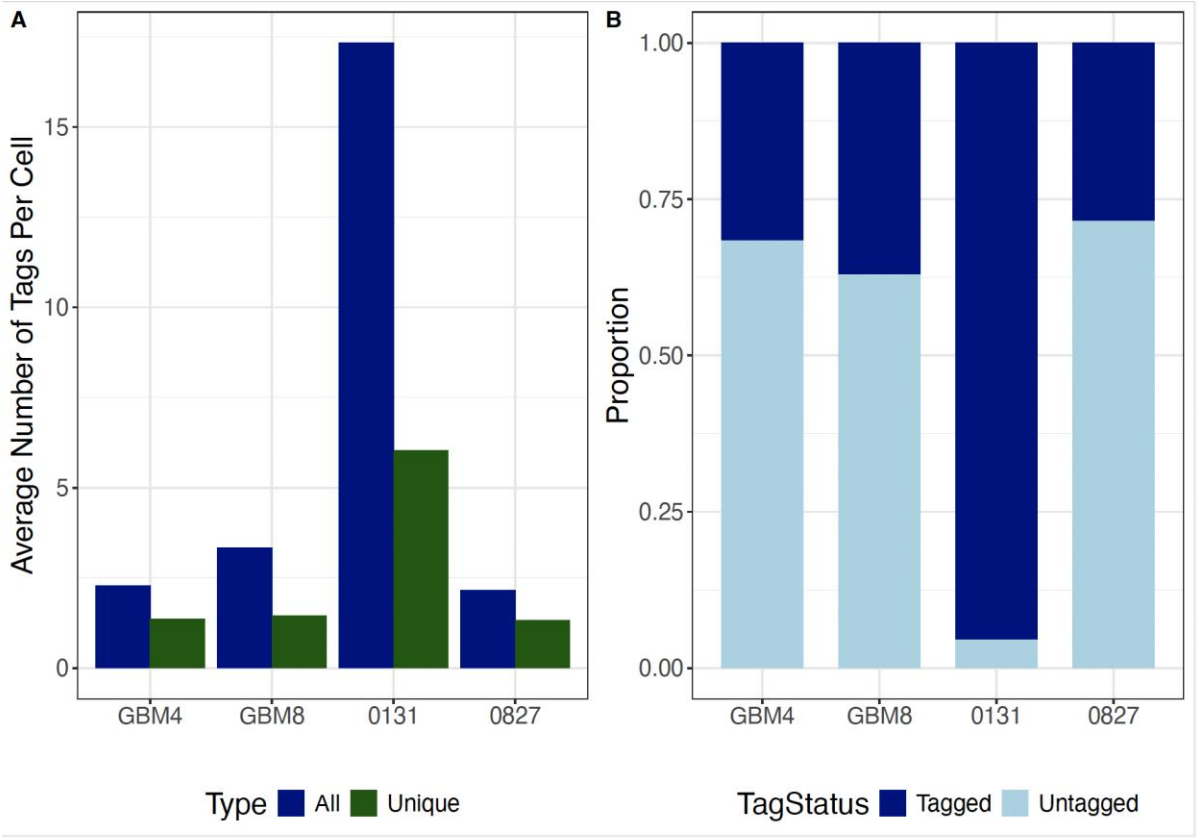


**Figure S13: CellTag labeling and distribution in GSC cultures.** (**A**) Average number of all CellTags and unique-sequence CellTags identified by amplicon sequencing in GSC cultures. (**B**) Proportion of single cells containing 1 or more CellTags, versus cells lacking identifiable CellTag sequences as detected and enumerated by scRNA sequencing.

**Figure S14: Total and unique CellTag barcode reads across GSC cultures, experimental arms and sampling time points.**

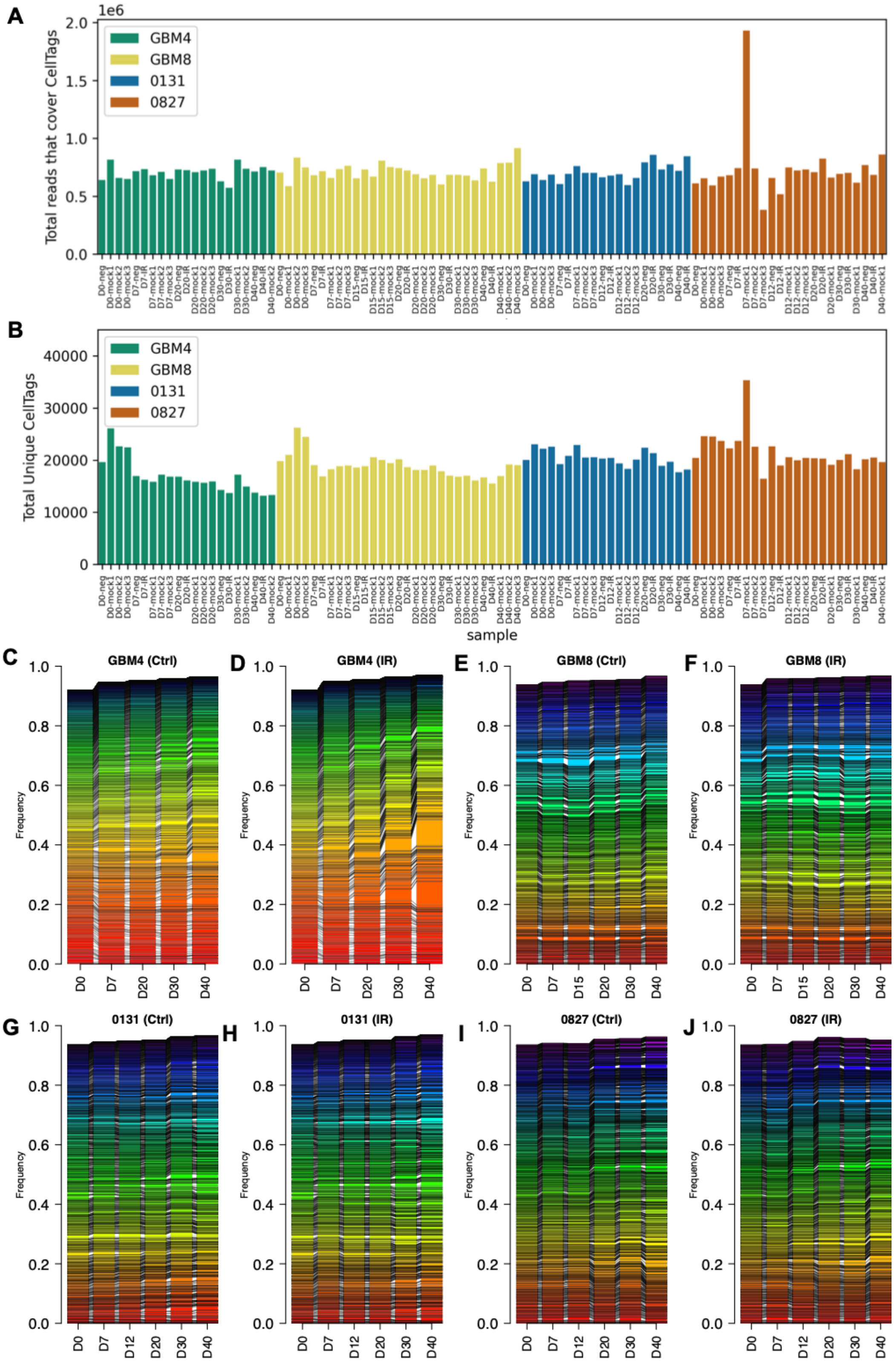

**Figure S14: Total and unique CellTag barcode reads across GSC cultures, experimental arms and sampling time points.** (A) Total reads per sample. (B) Total unique tags detected in each culture. (C-J) Cumulative read frequencies in control (Ctrl) and IR-treated (IR) GSC cultures sampled on specific days indicated along the X-axis over a 40 day time course sampling experiment. Unique-sequence tags with ≥ 10 reads at ≥ 1 time point were included.

**Figure S15: Single cell identification and relatedness identified from expressed, unique-sequence CellTag barcodes.**

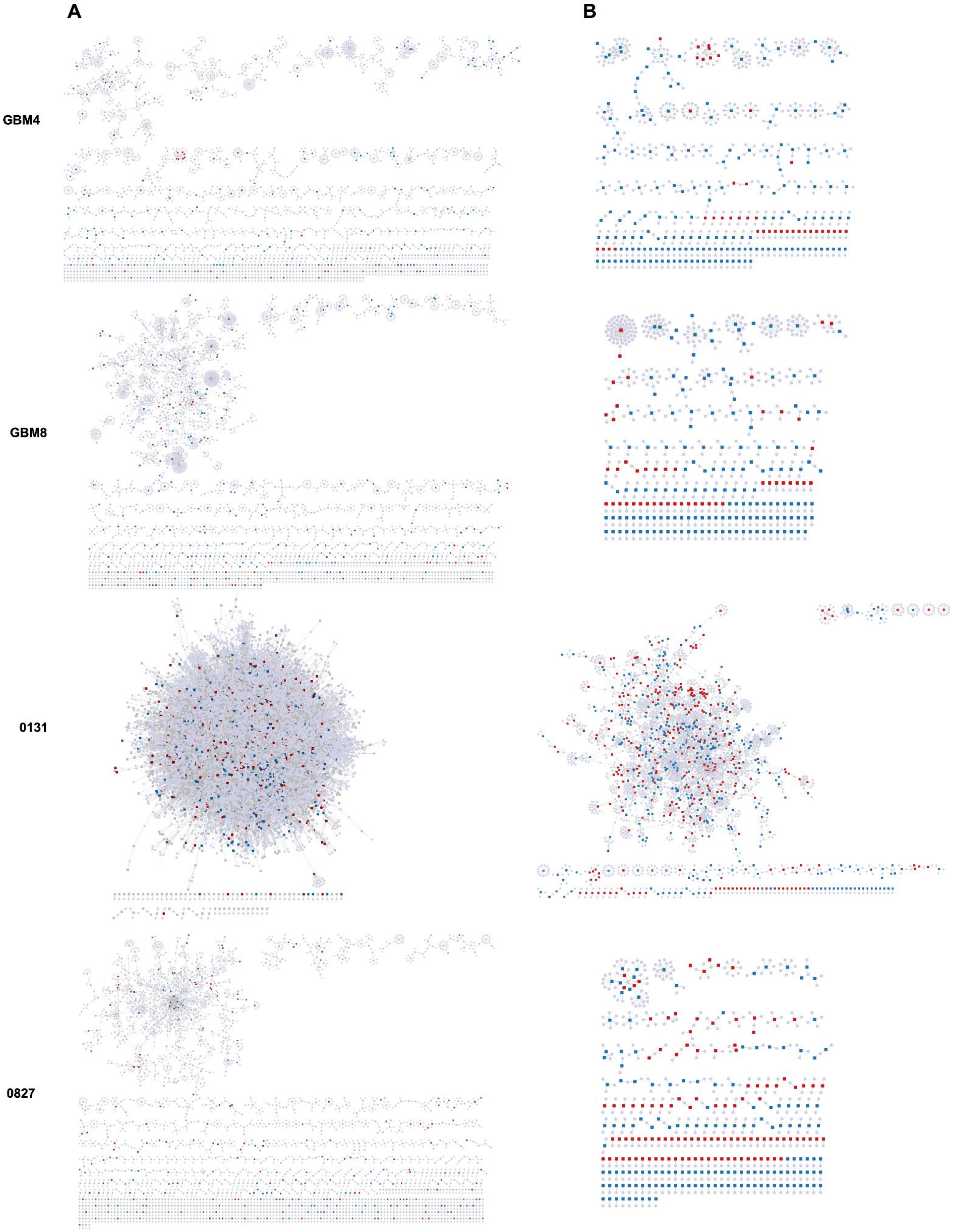

**Figure S15: Single cell identification and relatedness identified from expressed, unique-sequence CellTag barcodes.** (A) individual dots represent all single cells identified by cell-indexing barcodes included as part of the 10X single cell sequencing library prep at Day 1 of the time course sampling experiment depicted in Figure 5. Cells inter-related by 1 or more shared CellTag are indicated, as are enriched (red squares) or depleted (blue squares) tags identified by CellTag amplicon sequencing over the 40 day sampling time course. (B) topology plots for tagged single cells identified in (A) that were enriched (red squares) or depleted (blue squares) as a function of time over a 40 day sampling time course.

**Figure S16: Molecular barcode trajectories in untreated GSC cultures reveal tag and cellular enrichment/depletion over 40 days in cultures.**

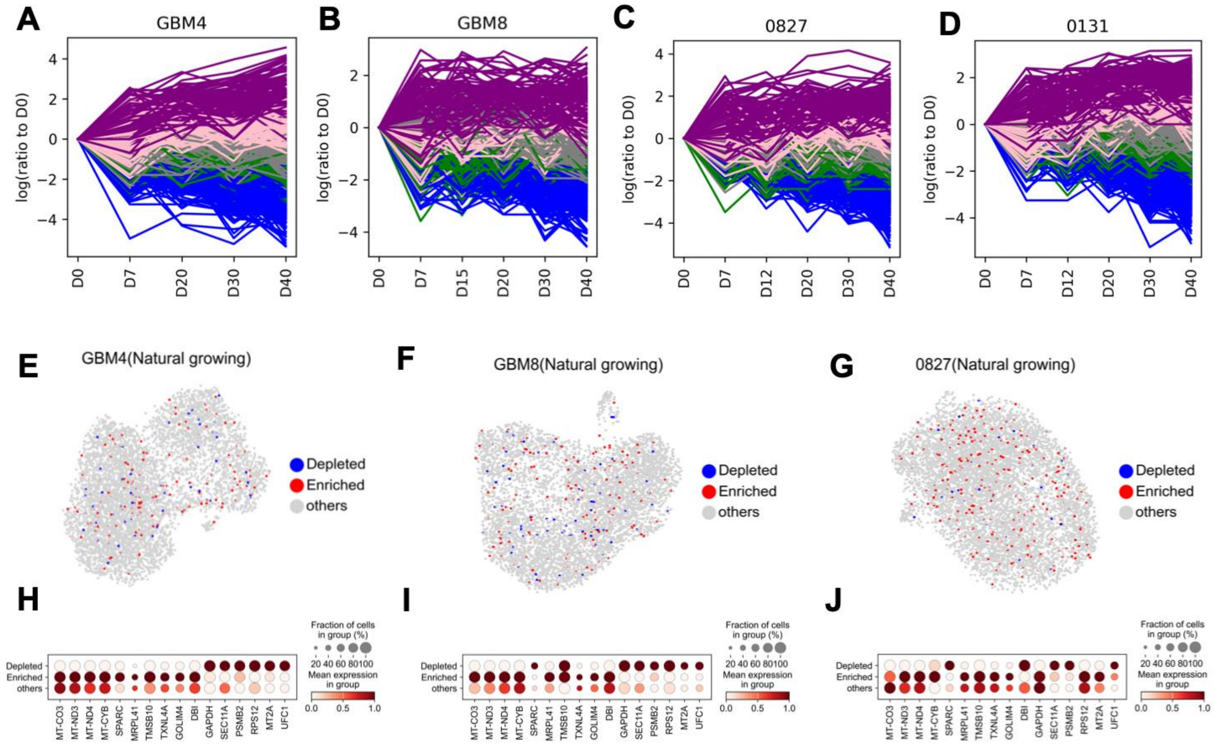


**Figure S16: Molecular barcode trajectories in untreated GSC cultures reveal tag and cellular enrichment/depletion over 40 days in cultures.** (A-D) Five groups of CellTag barcodes defined by K-means clustering in control cultures that were not IR-treated. Groups were identified on the basis of changes in frequency from Day 0 (D0) across 5 additional sampling time points to 40 days (D40). Clusters displayed trajectories ranging from clear depletion over 40 days (Cluster 0, blue) through no significant change (Cluster 2, gray) to clear enrichment (Cluster 4, purple) over 40 days. (E-G) Clearly depleted and enriched cells based on the CellTag content mapped onto D0 single cell UMAP plots for GBM4, GBM8 and 827. (H-J) Differentially expressed genes in enriched and depleted cell populations that exhibited differential expression in at least two or the three GBM4, GBM8 and 0827 cultures that were analyzed. Wilcoxon rank-sum test in the scanpy.tl.rank_genes_groups function is used to test the significance of differences. Threshold P values < 0.05 were used to identify differentially expressed genes.

**Figure S17: mtDNA variant trajectories over 40 days in control and IR-treated GSC cultures.**

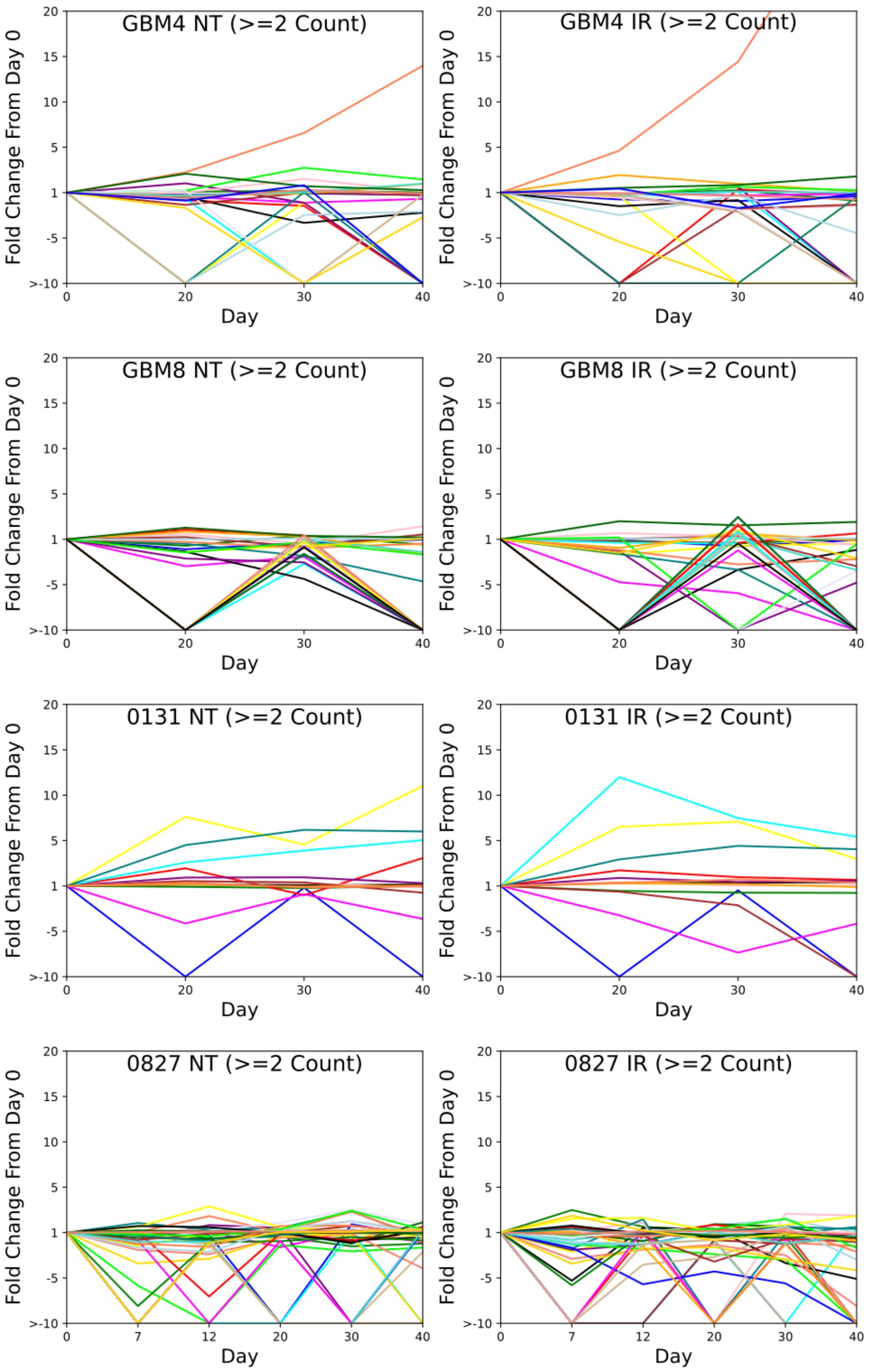

**Figure S17: mtDNA variant trajectories over 40 days in control and IR-treated GSC cultures.** Pre-existing (*i.e.* Day 0) mtDNA variants supported by at least two reads and are present in one or more subsequent time points can be tracked over time for both control (*left column*) and IR-treated (*right column*). Line colors are for the same variant in the respective control and IR-treated cultures: these reflect the highly dynamic nature of cells and subclones over time as identified by specific mtDNA variants. Most variants however displayed either no change over time or no change attributable to IR treatment.

#### Supplementary Tables

**Table S1: Short tandem DNA repeat allele length signatures unambiguously identify GSC cultures.** Cell line authentication STR profile signatures were defined using the CellCheck 9 Plus STR analysis panel (IDEXX BioAnalytics, Columbia, MO) which also was designed to detect *Mycoplasma* spp. infection and common cell line or species contaminants.

**Table S1: GSC culture STR authentication allele lengths**

| GSC | *AMEL* | *CSF1PO* | *D5S818* | *D7S820* | *D13S317* | *D16S317* | *THO* | *TPOX* | *vWA* |
|---|---|---|---|---|---|---|---|---|---|
| GBM4 | X | 11 | 10 | 10,12 | 12 | 8,11 | 9.3 | 8 | 17,18 |
| GBM8 | X | 11,13 | 12,13 | 8,10 | 12 | 11,12 | 9.2 | 8 | 18,19 |
| 0131 | X | 11 | 13 | 9,9.2 | 8 | 11 | 9.3 | 9 | 17 |
| 0827 | X | 12,13,(14) | 10,11,12 | (7.3),8,10 | 10,12,13 | 13,14 | 7 | 8,11 | 18,21 |

**Table S1 footnotes:** GSC culture 0827 STR fingerprinting revealed third alleles (indicated in parentheses) for two STR markers in expanded archival stocks versus initial cultures. These 3rd allele lengths are both for STR loci that reside on triplicated chromosomes in karyotypes of 0827 (see Figure S1).

**Table S2: Single nucleotide (SNV) and copy number (CNV) variants identified by GSC culture exome sequencing** (Excel spreadsheet)**.**

**Table S3: GSC culture Duplex mtDNA sequencing metrics** (Excel spreadsheet)**.**

**Table S4:  GSC culture-specific mtDNA variants identified by Duplex mtDNA sequencing** (Excel spreadsheet)**.**

**Table S5: Cell Ranger summary statistics from GSC culture scRNAseq analyses.**

| Cell Ranger count pipeline outputs for GBM4, GBM8, 0131 and 0827 cultures | | | | |
|---|---|---|---|---|
| | **GBM4** | **GBM8** | **0131** | **0827** |
| **Estimated Number of Cells** | 5,973 | 5,906 | 7,502 | 7,400 |
| **Mean Reads per Cell** | 23,875 | 23,720 | 20,024 | 14,909 |
| **Median Genes per Cell** | 2,653 | 2,648 | 2,695 | 2,059 |
| **Number of Reads** | 142,606,568 | 140,091,044 | 150,225,708 | 110,328,090 |
| **Valid Barcodes** | 97.6% | 97.5% | 97.2% | 97.2% |
| **Sequencing Saturation** | 28.8% | 29.6% | 30.0% | 25.3% |
| **Q30 Bases in Barcode** | 94.9% | 95.0% | 94.9% | 94.8% |
| **Q30 Bases in RNA Read** | 92.7% | 92.5% | 92.6% | 91.9% |
| **Q30 Bases in UMI** | 94.5% | 94.4% | 94.4% | 94.3% |
| **Reads Mapped to Genome** | 97.1% | 97.7% | 97.3% | 96.6% |
| **Reads Mapped Confidently to Genome** | 93.8% | 94.4% | 94.7% | 92.9% |
| **Reads Mapped Confidently to Intergenic Regions** | 6.1% | 6.2% | 5.6% | 5.7% |
| **Reads Mapped Confidently to Intronic Regions** | 27.8% | 29.8% | 29.3% | 20.8% |
| **Reads Mapped Confidently to Exonic Regions** | 59.9% | 58.4% | 59.8% | 66.3% |
| **Reads Mapped Confidently to Transcriptome** | 56.2% | 54.7% | 56.0% | 61.8% |
| **Reads Mapped Antisense to Gene** | 1.4% | 1.5% | 1.6% | 1.3% |
| **Fraction Reads in Cells** | 93.9% | 94.3% | 96.0% | 92.0% |
| **Total Genes Detected** | 21,345 | 21,359 | 21,945 | 19,859 |
| **Median UMI Counts per Cell** | 8,309 | 7,646 | 6,637 | 5,802 |

**Table S6: Distribution of gene expression-defined GBM subtypes from single cell gene expression profiling data.**

| Culture | AC | MES | NPC | OPC | Undetermined |
|---|---|---|---|---|---|
| **GBM4** | 2.00% | 27.30% | 9.60% | 4.40% | 56.70% |
| **GBM8** | 0.80% | 0.60% | 25.80% | 14.30% | 58.40% |
| **0131** | 2.80% | 63.80% | 1.20% | 0.30% | 31.90% |
| **0827** | 6.20% | 17.00% | 16.70% | 0.70% | 59.40% |

**Table S7: Expressed mtDNA variants and expressed CellTags identified in GSC cells by scRNAseq UMI indexing.**

**Table S7. Number of expressed mtDNA variants and CellTags identified in GSC cultures by scRNAseq UMI indexing**

| | Total number of cells identified by 10X UMI | UMI cells with expressed CellTag | UMI cells with expressed mtDNA variant | UMI cells identified by expressed CellTag and mtDNA variant | Number of unique CellTag bar codes | CellTag unique variant counts | CellTag variant count range | mtDNA unique variant counts | mtDNA variant count range |
|---|---|---|---|---|---|---|---|---|---|
| **GBM4** | 5971 | 1887 | 8 | 3 | 612 | 1 | [1-6] | 7 | [1-1] |
| **GBM8** | 5900 | 2187 | 62 | 29 | 640 | 0 | [13-19] | 45 | [1-1] |
| **131** | 7495 | 7153 | 14 | 12 | 1306 | 0 | [5-35] | 52 | [1-2] |
| **827** | 7396 | 2108 | 7363 | 2106 | 837 | 0 | [24-1408] | 9 | [1-11] |

**Table S8: Gene sets for transcription functional characterizations.**

| Geneset | Descriptive Summary | Source |
|---|---|---|
| Cell Surface Markers | Literature curated cell surface markers | Wakimoto et al. 2009 PMID: 19351838/PMCID: PMC2785462; Son et al. 2009 PMID: 19427293/ PMCID: PMC7227614; Kenney-Herbert et al. 2015 PMID: 26019225/PMCID: PMC4479614; Brown et al. 2015 PMID: 25749043/ PMCID: PMC4467436 |
| Cell Cycle Genes | Signature genes for G1/S and G2/M cell cycle phases | Table S2 of Neftel et. al. Cell 2019 (PMID: 31327527) |
| Stemness Signature | List of Published Stem Cell Markers (Healthy and Cancer). Geneset includes Yamanaka factors, known normal stem cell markers and GBM-specific cancer stem cell markers | Table S2 from Malta et. al. Cell 2018 (PMID: 29625051) |
| MSigDB Hallmark 50 | Genes that display coordinate expression and represent well-defined biological processes. | MSigDB Hallmark Genesets (PMID: 26771021) |
| Wang GBM Intrinsic Subtypes | Signature genes that defined the mesenchymal, proneural, classical GBM intrinsic subtypes as defined in TCGA | Table S1 of Wang et. al. Cancer Cell 2017 (PMID: 28697342) |
| Neftel Cell State | MES1-like, MES2-like, NPC1-like, NPC2-like, AC-like, OPC-like malignant state markers. Meta-modules defined as genes with average log-ratios > 2 and restricted to the top 50 genes with highest log-ratios for signature, with genes are listed descending by average log-ratios. | Table S2 of Neftel et. al. Cell 2019 (PMID: 31327527) |
| Garafano Metabolic Subtypes | Signature genes for the 4 GBM subtypes: Glycolytic/plurimetabolic (GPM), mitochondrial (MTC), neuronal (NEU), proliferative/progenitor (PPR) | Supplementary Table 6J of Garofano et. al. Nat Cancer 2021 (PMID: 33681822) |
| Invasivity Gene Signature | Gene Signatures Associated with Tissue States | Table S7 of Venkataramani et. al. Cell 2022 (PMID: 35914528) |
| AMPARs | α-amino-3-hydroxy-5-methyl-4-isoxazolepropionic acid receptor-type glutamate receptors (AMPARs) | STAR Methods 'Definition of single-cell invasivity signature and AMPAR scores' from Venkataramani et. al. Cell 2022 (PMID: 35914528) |
| Connectivity Upregulated | Gene signature derived from scRNA-Seq data that defines genes up or down- regulated in highly connected cells | Supplemental Data 1 from Hai et. al. Nat Commun 2024 (PMID: 38320988) |
| Connectivity Downregulated | | |

**Table S9: CellTag reads for time course sampling experiment** (Excel spreadsheet)**.**

**Table S10: Gene set enrichment statistics for bulk RNA and scRNA sequencing data analyses** (Excel spreadsheet)**.**

**Table S11: Protein and phosphoprotein abundances and statistics** (Excel spreadsheet)**.**
